# Selection on ancestral genetic variation fuels parallel ecotype formation in bottlenose dolphins

**DOI:** 10.1101/2020.10.05.325159

**Authors:** M. Louis, M. Galimberti, F. Archer, S. Berrow, A. Brownlow, R. Fallon, M. Nykänen, J. O’Brien, K. M. Roberston, P. E. Rosel, B. Simon-Bouhet, D. Wegmann, M.C. Fontaine, A.D. Foote, O.E. Gaggiotti

**Affiliations:** Scottish Oceans Institute, University of St Andrews, East Sands, St Andrews KY16 8LB, Scotland, UK; Centre d’Etudes Biologiques de Chize, Universite de La Rochelle 17000 La Rochelle, France; Groningen Institute for Evolutionary Life Sciences (GELIFES), University of Groningen, PO Box 11103 CC, Groningen, The Netherlands; Globe Institute, University of Copenhagen, Øster Voldgade 5, 1350 Copenhagen, Denmark; Department of Biology and Biochemistry, University of Fribourg, Fribourg, Switzerland; Swiss Institute of Bioinformatics, Fribourg, Switzerland; Marine Mammal and Turtle Division, Southwest Fisheries Science Center, NOAA, 8901 La Jolla Shores Drive, La Jolla, CA 92037, USA; Irish Whale and Dolphin Group, Kilrush, Co Clare, Ireland; Marine and Freshwater Research Centre, Department of Natural Sciences, School of Science and Computing, Galway-Mayo Institute of Technology, Dublin Road, H91 T8NW Galway, Ireland; Scottish Marine Animal Stranding Scheme, SRUC Northern Faculty, An Lòchran, Inverness Campus, IV2 5NA,UK; School of Medicine, University of St Andrews, North HaughSt Andrews, Fife, KY16 9TF, Scotland, UK; University College Cork, Cork, Ireland; NOAA Fisheries, Southeast Fisheries Science Center, NOAA, 646 Cajundome Boulevard, Lafayette, LA 70506, USA; MIVEGEC (Université de Montpellier, CNRS 5290, IRD 229) et Centre de Recherche en Écologie et Évolution de la Santé (CREES), Institut de Recherche pour le Développement (IRD), F-34394, Montpellier, France; Molecular Ecology Fisheries Genetics Lab, School of Biological Sciences, Bangor University, Bangor, UK; Department of Natural History, Norwegian University of Science and Technology (NTNU), NO-7491 Trondheim, Norway

## Abstract

What are the mechanisms that allow species to extend their ranges and adapt to the novel environmental conditions they find in the newly available habitat? The study of parallel adaptation of pairs of populations to similar environments can provide great insights into this question. Here, we test for parallel evolution driven by niche specialization in a highly social marine mammal, the common bottlenose dolphin, *Tursiops truncatus*, and investigate the origins of the genetic variation driving local adaptation. Coastal ecotypes of common bottlenose dolphins have recurrently emerged in multiple regions of the world from pelagic ecotype populations, when novel habitat became available. Analyzing the whole genomes of 57 individuals using comparative population genomics approaches, we found that coastal ecotype evolution was relatively independent between the Atlantic and Pacific, but related between different regions within the Atlantic. We show that parallel adaptation to coastal habitat was facilitated by repeated selection on ancient alleles present as standing genetic variation in the pelagic populations. Genes under parallel adaptation to coastal habitats have roles in cognitive abilities and feeding. Therefore, parallel adaptation in long-lived social species may be driven by a combination of ecological opportunities, selection acting on ancient variants, and stable behavioural transmission of ecological specialisations. Tried and tested genetic variation that has been subject to repeated bouts of selection, may promote linked adaptive variants with minimal pleiotropic effects, thereby facilitating their persistence at low frequency in source populations and enabling parallel evolution.

---

Understanding the processes that allow species to extend their ranges and adapt to novel environments is a long-standing question in biology, which interest now extends well beyond this disciplinary field due to the potential effect of global change on species ranges. The colonisation of novel environments may result in new selective pressures on individuals and promote local adaptation (1). However, linking genetic divergence to local adaptation is particularly challenging as genetic differentiation may also arise due to demographic history (2), or other selective processes such as background selection (3). Replicate adaptation of different populations to similar environments is often considered strong evidence of the repeated action of natural selection (4). Hence, we can study parallel evolution to gain insights into the mechanisms driving genetic variation and adaptation.

Iconic examples of parallel evolution include adaptation to similar environments, i.e. repeated independent colonisation of freshwater environments from marine habitats in threespine sticklebacks, *Gasterosteus aculeatu*s (5), parallel adaptation to the same host species in stick insects, *Timema cristinae* (6), high altitude adaptation in multiple human, *Homo sapiens*, populations (7), and different light conditions in cichlid fish (8), or similar responses to comparable stressors (e.g. virus (9) or pollution (10). Our understanding of the mechanisms involved has recently shifted from a binary view of repeated *vs*. idiosyncratic processes to a continuum ranging from parallel to non-parallel (6, 11–14).

Parallel evolution may occur rapidly, if the genetic substrate which selection acts upon was already segregating in the ancestral population (i.e. standing genetic variation, SGV (5, 6, 9–11, 15, 16)) as balanced polymorphisms (17), or introgressed from a locally adapted outgroup (18). Alleles present as SGV may have been selected in past environments, potentially increasing their chances to be the target of natural selection (15). Recent studies have highlighted that the origin of the alleles that enable populations to recurrently adapt to similar environments may be much older than the divergence of the populations themselves (16, 19, 20). For example, the reservoir of alleles in marine populations of threespine sticklebacks, that have been recurrently selected after freshwater colonisation during the past 12,000 years, has been segregating for millions of years (20).

With the rare exception of humans (7, 21), reported cases of parallel evolution almost exclusively involve relatively short-lived species (5, 6, 8–10, 16, 22). In long-lived species, such as large mammals, long generation time, low fecundity and sometimes small effective population size may hamper rapid local adaptation, especially from de novo mutation (15, 23). In humans, parallel adaptation likely resulted from cultural innovations, such as lactase persistence in different pastoral populations as a result of animal domestication (21), or living at high altitude (7). In other long-lifespan social mammals, stable social transmission of learned behaviours such as foraging strategies or habitat preferences may also facilitate the evolution of local adaptation, although examples are scarce (but see killer whales, *Orcinus orca* (24)).

Here, we tested for parallel evolution driven by ecological niche specialization in a highly social marine mammal, the common bottlenose dolphin, which has a worldwide temperate and tropical distribution. Two ecotypes of common bottlenose dolphins (coastal and pelagic, also called offshore) have recurrently formed in multiple regions of the world (25–29). Coastal populations were suggested to have been founded from pelagic source populations (25–27, 30). They are thus an excellent study system to test whether parallel evolution occurred and involved the same molecular processes during the repeated colonization of coastal habitat. Throughout their range, coastal populations have different diets compared to pelagic populations (31–33), and display phenotypic traits adapted to coastal waters, in particular for feeding (33–35). Coastal populations in distinct regions can share some morphological traits like larger teeth, rostra, and internal nares when compared to pelagic populations. They can also show some unique traits and adaptations such as North-West Atlantic (NWA) coastal bottlenose dolphins which are smaller than their pelagic counterparts, while in other regions the pattern is reversed or there are no discernable differences (31, 33–35). Coastal populations tend to show strong site fidelity and reduced dispersal (27, 29, 36), and foraging ecology is transmitted both vertically (from mother to calves) and socially (from conspecifics in the social groups) (37, 38). Resident behaviour and habitat-specific foraging traditions would be expected to facilitate the evolution of local adaptation.

The aim of our study was to identify the mode of evolution at the molecular level underlying repeated divergence in pelagic and coastal common bottlenose dolphins. We first identified their population structure and demographic history. We showed the pelagic and coastal ecotype pairs have evolved independently between the Atlantic and the Pacific but have a partially shared history in the Atlantic. Then, we characterised parallel patterns of selection to coastal habitat across the genome, and identified genes under parallel evolution potentially involved in cognitive abilities and feeding.

## Results and discussion

### Genetic structure

We analysed nuclear genomes (7.54 X ± 1.52 after quality filtering) of 57 common bottlenose dolphins (Figure 1, Supplementary table 1), including ten coastal and ten pelagic individuals from the North-East Atlantic (NEA), ten pelagic and seven coastal individuals from the North-West Atlantic (NWA) and nine coastal and eleven pelagic individuals from California, North-East Pacific (NEP).

**Figure 1.**
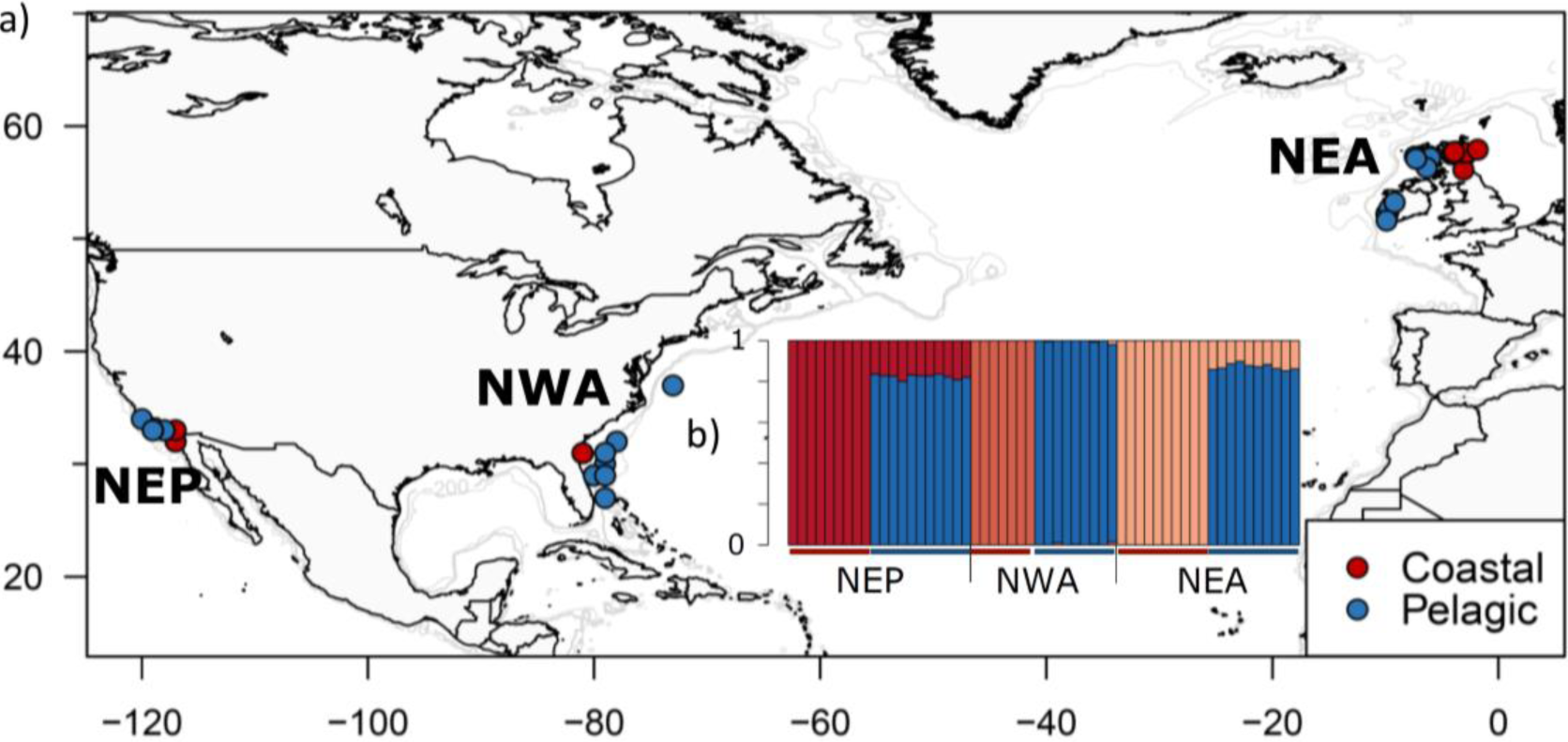
Sampling location and genetic ancestry of each coastal and pelagic common bottlenose dolphin populations. a) Map of sample locations of the common bottlenose dolphin ecotypes, in the North-East Atlantic (NEA), North-West Atlantic (NWA) and North-East Pacific (NEP) and b) ancestry proportions for each of the 57 individuals inferred in NGSAdmix(40) for a number of clusters, K=4, identified as the highest level of structure using the Evanno method(75).

The genetic structure obtained from a principal component analysis (PCA) (39) and the individual-based ancestry and clustering analysis of NGSAdmix (40) based on a set of 798,572 unlinked high quality SNPs indicated that the samples assigned *a priori* to a population clustered together. The analyses showed two major axes of differentiation: Atlantic *vs*. Pacific, and pelagic *vs*. coastal (Figure 1, Supplementary figures 1-3, pairwise *F*_ST_ in Supplementary table 2). The three pelagic populations were more closely related to each other, even when in different ocean basins, than any of them were to a parapatric coastal population. In contrast, the three coastal populations were very differentiated from each other (Figure 1, Supplementary figures 1-3, pairwise *F*_ST_ in Supplementary table 2). These patterns of differentiation suggest that the coastal populations resulted from several founding events.

### Demographic history

To investigate this hypothesized founder history of coastal populations, we used coalescent-based estimates of effective population size on putatively neutral regions identified by Flink (41), which may reflect either changes in population size or population structure (42). We found that pelagic populations experienced demographic expansions followed by a period of more stable *N*_e_ than the coastal populations (Figure 2a, Supplementary figures 4, 5a-b). Population expansion at the start of the last glacial period may reflect increased connectivity, rather than an increase in *N*_e_, as suitable habitat became scarce (30). All coastal populations went through a bottleneck followed by an expansion, which indicates that founder effects were associated with coastal ecotype formation. Reduced nucleotide diversity, Watterson’s Theta, and consequently positive Tajima’s D estimates (Figure 2c, Supplementary figures 6a-b and 7) also indicate that the coastal populations have experienced bottlenecks and suggest they were derived from the pelagic populations. Access to novel previously ice covered shallow coastal habitats during past climate change at the end of the LGM in the NEA (30, 31), or during warm interstadials created opportunity for ecological differentiation. Coastal habitats provide a mosaic of environments and different and potentially more stable food resources (31, 43).

**Figure 2.**
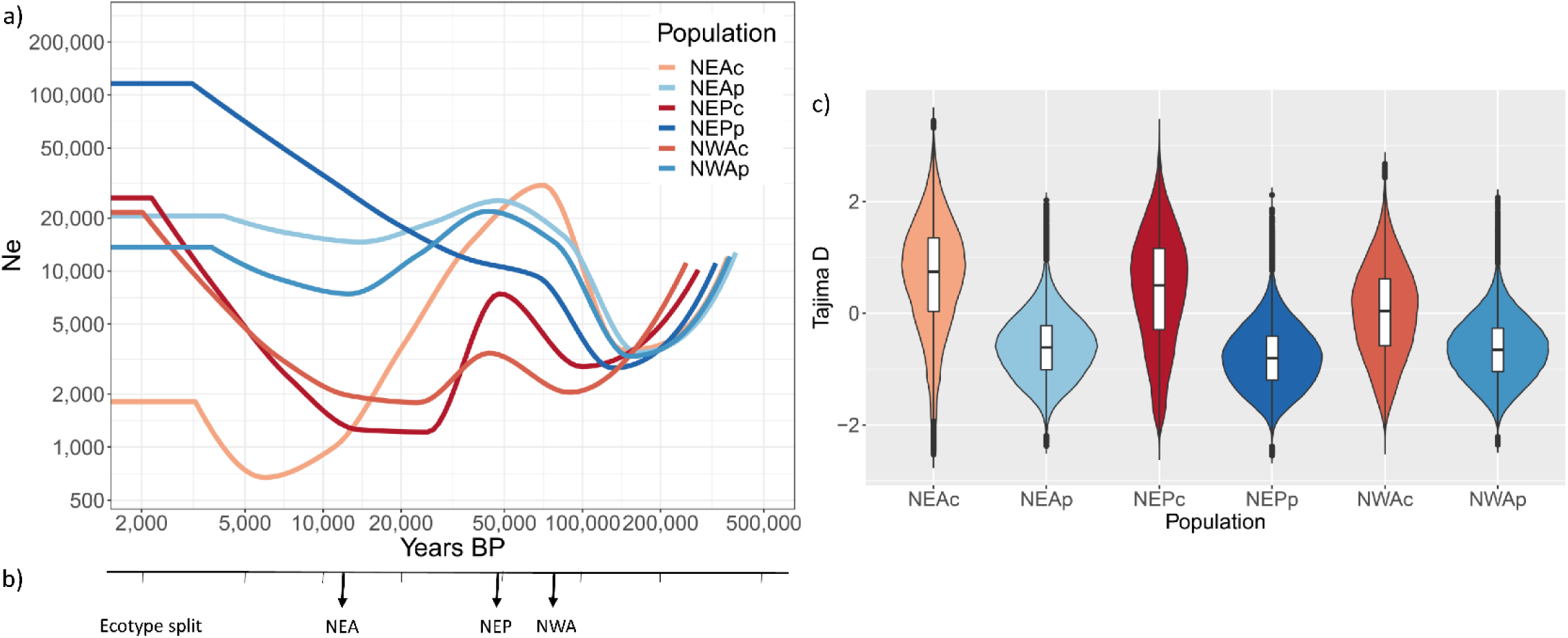
Demographic history of common bottlenose dolphin populations. a) Changes in effective population size through time inferred for each common bottlenose dolphin population using SMC++ using a mutation rate of 2.56e-8 substitution per nucleotide per generation(81) and a generation time of 21.1 years(79). b) Split time between ecotypes in each region estimated using SMC++. Populations are North-East Atlantic coastal (NEAc), North-East Atlantic pelagic (NEAp), North-East Pacific coastal (NEPc), North-East Pacific pelagic (NEPp), North-West Atlantic coastal (NWAc) and North-West Atlantic pelagic (NWAp). c) Tajima D estimated for each population, the **violin plots** indicate the kernel probability density of the data, the box indicates the interquartile range and the horizontal marker the median of the data.

**Figure 3.**
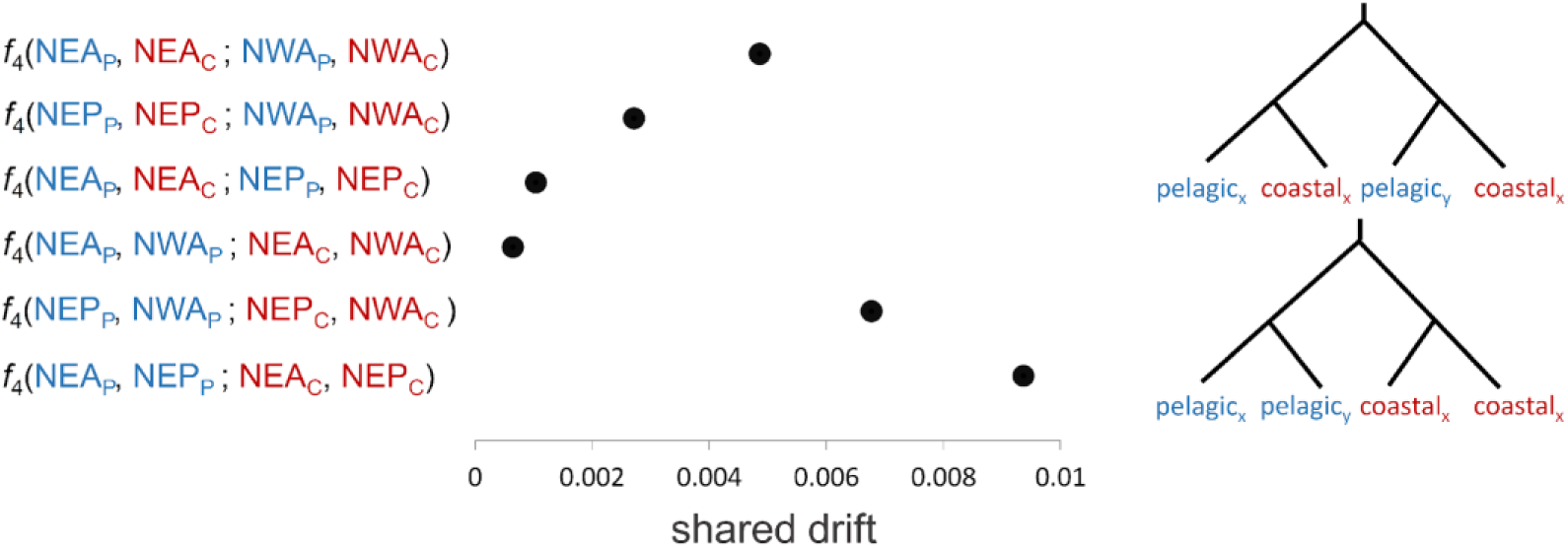
Admixture among populations of common bottlenose dolphins. *F*_4_-statistics of the form *F*_4_(pelagic_x_, coastal_x_; pelagic_y_, coastal_y_) or *F*_4_(pelagic_x_, pelagic_y_; coastal_x_, coastal_y_). All the SE estimations are less than 1e-4 and all *F* _4_-statistics were significant based on Z-scores greater than 3, which is the equivalent of a significance of *P* <0.0026

**Figure 4.**
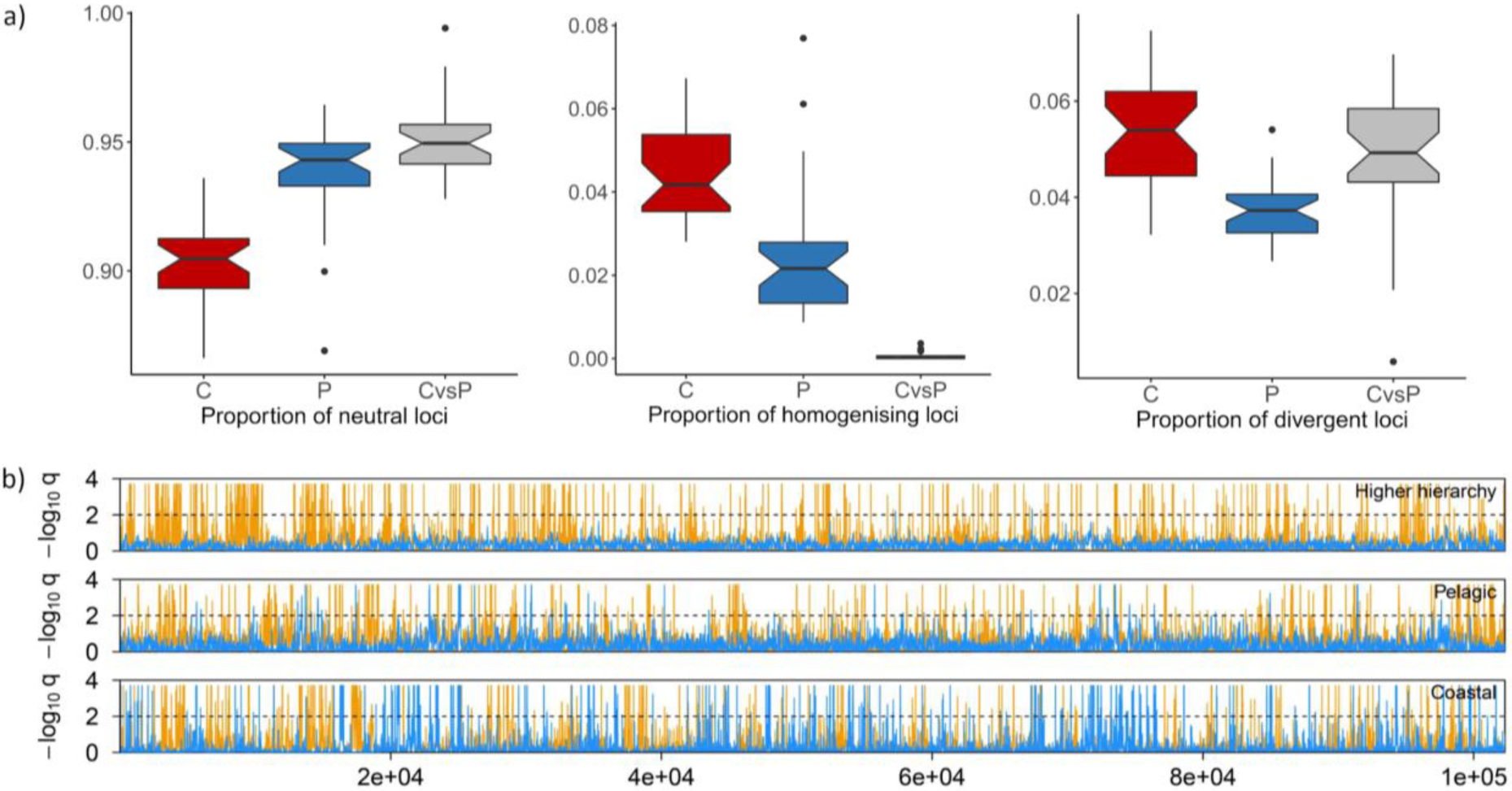
Patterns of selection within and between common bottlenose dolphin ecotypes. a) Boxplots of the genomic patterns of selection within coastal (C) and pelagic (P) ecotypes and between the two ecotypes (CvsP), proportion of neutral, homogenizing and divergent loci. b) Patterns of selection (divergent: yellow, homogenising: blue) inferred using Flink from one scaffold grouping between coastal and pelagic population (top panel), among pelagic populations (middle panel) and among coastal populations (lower panel) for one scaffold ensemble. The y-axis indicates the locus-specific FDR for divergent (orange) and homogenising (blue) selection, respectively. The black dashed line shows the 1% FDR threshold, above which we consider the locus under selection.

**Figure 5.**
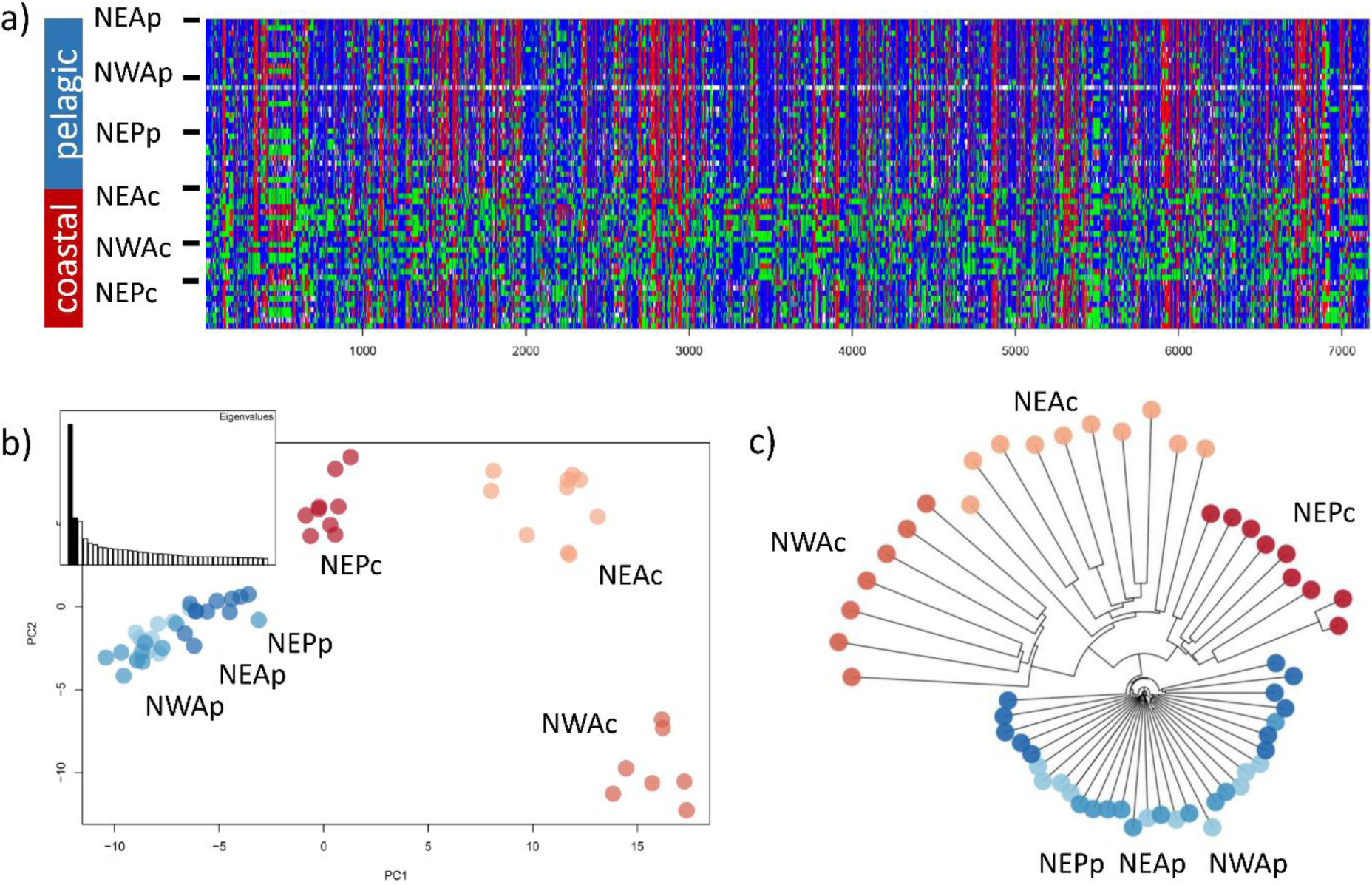
Patterns of genetic variation of the 7,165 SNPs under parallel selection to coastal habitat, i.e. under both homogenising selection among coastal population and divergent selection between ecotypes. These SNPs are scattered across the genome. a) Plot of the homozygote reference genotypes in blue, heterozygote in green and homozygote for the alternated allele in red. b) Principal component analysis and (c) Neighbor-joining distance tree showing the genetic structure of the common bottlenose dolphin samples for this particular SNP set.

### Evolutionary relationships among ecotypes

To establish if our three geographic pairs of coastal and pelagic ecotypes represent independent ecological divergence events, we reconstructed their demographic histories using approaches that estimate covariance of drift from allele frequencies. We first explored the demographic history and potential admixture events using TreeMix (44), which supported the independent split of the Pacific and Atlantic coastal populations (Supplementary figures 9 and 10, Supplementary text). In contrast, the two coastal populations in the eastern and western North Atlantic were closely related. The TreeMix results also indicated the contribution of an outgroup, the strictly coastal sister species, the Indo-Pacific bottlenose dolphin, *Tursiops aduncus*, used to root the Treemix tree, to the genetic composition of the NWA coastal population. However, the addition of this migration edge resulted in incongruence among *T. truncatus* populations, indicating the admixture graph was still not a good fit for the data. D-statistic tests further reject direct introgression between the NWAc population and *T. aduncus* (Supplementary table 6). This outgroup may not be *T. aduncus*, but could be a lineage with ancestry shared with *T. aduncus*. Alternatively, shared ancestral polymorphisms that differentially segregate in coastal *T. truncatus* and *T. aduncus* lineages may underlie this pattern, see Discussion below.

Given the incongruence of the TreeMix graph to the data, we further tested whether geographic pairs of pelagic and coastal ecotypes had evolved independently by estimating the less parameterized *F*_4_-statistics (45, 46) of the form (pelagic_x_, coastal_x_; pelagic_y_, coastal_y_), where x and y represent different geographic regions. In contrast to the expectation in case of independent colonisations, *F*_4_-statistics were significantly positive (Figure 3). The strongest signal of non-independence was observed between the two Atlantic ecotype pairs, which suggests a partially shared evolutionary history of parallel coastal and pelagic ecotypic divergences within the North Atlantic. We also calculated *F*_4_-statistics comparing drift between the same ecotypes from different geographic regions, *F*_4_(pelagic_x_, pelagic_y_; coastal_x_, coastal_y_). Tests of this form were all significantly positive, and *F*_4_(NEA_p_,NWA_p_;NEA_c_,NWA_c_) was the lowest of all and, importantly, lower than *F*_4_(NEA_p_,NEA_c_; NWA_p_,NWA_c_) suggesting that the two Atlantic coastal populations may be derived from the same ancestral pelagic population. Thus, overall, *F*_4_ results are consistent with the TreeMix results and suggest that coastal habitat colonisations by common bottlenose dolphins in the Atlantic are not fully independent. On the other hand, colonisation of coastal habitats in the East Pacific was distinct from founder events in the Atlantic, but nevertheless involved some admixture with Atlantic (or closely related) populations.

We estimated divergence time between the two ecotypes within each region using SMC++ on putatively neutral regions. The oldest divergence between coastal and pelagic ecotype occurred in the NWA (around 80,000 yBP), and the youngest was a post-glacial divergence in the NEA (Figure 2b, Supplementary figures 11a-b). Divergence time between the two coastal populations in the North Atlantic was estimated around 50,000-70,000 yBP (Supplementary figures 12a-b). This rather old divergence is inconsistent with their position in the TreeMix tree, which indicates they are closer to each other than the NEA coastal population is to the NEA pelagic population.

One mechanism to explain differences in the chronology of divergence between population pairs inferred by different analyses may be considering the genome as a mosaic of different evolutionary histories. For example, tracts of older ancestry components would be expected to have accumulated more mutations, which segregate among populations. To test for that, we search for tracts of dense private mutations (47) segregating in each coastal individual relative to the allopatric pelagic populations (Supplementary text and Supplementary figure 13). The highest density of private mutations was found in the NWAc dolphins (Supplementary tables 3-4). Three to five times more tracts of dense private mutations were shared between the coastal populations of the NEA and NWA than between the coastal populations of the NEA and NEP and of the NWA and NEP, in line with their partially shared ancestry inferred from the *F*_4_-statistics and TreeMix (Supplementary table 5). We hereafter refer to such tracts as ‘ancient’ given their older TMRCA (0.6 to 2.3 *vs*. 0.09-0.4 million years old for the rest of the genome).

The highest density of ancient tracts in the NWAc individuals are consistent with the TreeMix results indicating the contribution of an outgroup to the genetic composition of the NWA coastal dolphins (Supplementary figures 9-10). Evidence of ancestral introgression from extinct or unsampled (i.e. ghost) populations or species has recently been found in other marine species (e.g killer whales and sea bass (48, 49)). However, these ‘ancient’ tracts do not need to be directly introgressed, but may rather have been retained as balanced polymorphisms (17). The presence of such ancient tracts may also explain the basal position of the NWA coastal population in a phylogenetic substitution-based inference that assumes a simple bifurcating branching process using RAD-sequencing data (50). The divergence dates of *T. aduncus* and *T. truncatus* estimated by Moura et al. 2020 (50) and McGowen et al. 2020 (51) are close to the TMRCA of ancient tracts found in NWAc (1.0-2.3 million years old), after correcting for the different mutation rates used between studies.

LocalPCA (52), a method that describes heterogeneity in patterns of relatedness among populations, further confirmed that our dolphin genomes were comprised of regions with different evolutionary histories. The three major evolutionary relationships (on PCs 1 and 2) include i) the pattern expected under the scenario of each coastal population originating from the pelagic population in the same region, ii) the pattern where the NEA and NWA coastal populations were more closely related than expected under independent ecotype splits on each side of the Atlantic (Supplementary figures 14-15), as supported by the *F*_4_-statistics and TreeMix (Figure 3, Supplementary Figure 9) and iii) a pattern where the NWA coastal population was closer to the pelagic populations. Furthermore, on PCs 3 and 4, coastal populations from the Atlantic and Pacific clustered together and likewise for the pelagic populations, suggesting parallel ecotype-based processes (Supplementary figures 14-15).

### Mechanisms of parallel evolution to coastal habitat

To test whether the above results can be interpreted as parallel selection associated with coastal habitat, we used Flink (41), an extension of BayeScan (53) that takes linkage among loci into account. We considered ecotype (coastal *vs*. pelagic) as the top hierarchical level, followed by the three populations within each group. Our analyses show striking differences in patterns of inferred selection involving mainly divergent selection between coastal and pelagic ecotypes (higher hierarchy), and mainly homogenising selection (less genetic differentiation than expected under neutrality) among coastal populations (Figure 4a-b, supplementary text and figures 16-17a). Homogenising selection implies that the same alleles have been repeatedly selected in the different coastal populations. We acknowledge that divergent selection within and between ecotypes may be inflated by false positives associated with bottlenecks in the founding of coastal populations. Plots of all neutral and selected raw genotypes show more variability in pelagic populations and more fixed alleles in the coastal populations, suggesting coastal populations were derived from repeated colonisation utilising genetic variation present in pelagic populations (Supplementary figures 16-17b).

We then explored the possible origins of the variants under selection. Our results show that most of them were polymorphic, i.e. present as SGV, in the pelagic populations. Ninety-six percent of the 89,796 outlier SNPs under homogenising selection and 72% of the 89,663 loci under diverging selection were polymorphic in the pelagic populations (Supplementary text and supplementary figures 18-20). We consider the 7,165 SNPs being under both homogenising selection among coastal populations and diverging selection between ecotypes as putative loci underlying parallel evolution to coastal habitats (Figure 5a-c, Supplementary text) and focus on those variants in the rest of our study. Most (81%) of those SNPs were polymorphic in the pelagic populations. On a PCA and unrooted NJ tree based on these 7,165 SNPs, the populations clustered by ecotype, further supporting their having evolved under parallel selection, and potentially having a role in habitat adaptation (Figure 5b-c). Additionally, when including the *T. aduncus* individual, which is coastal, in the PCA and NJ tree those SNPs showed a pattern of greater coalescence between *T. aduncus* and *T. truncatus* coastal individuals, than between *T. truncatus* coastal and *T. truncatus* pelagic individuals, further supporting their implication in coastal habitat adaptation (Supplementary figure 21). This topology is discordant with that of most of the genome, but concordant with the covariance in a subset of allele frequencies detected by TreeMix.

We found that a large proportion (66%) of the SNPs under parallel selection in coastal populations were found in ancient (0.6 to 2.3 million years old) tracts. In contrast, only an average of 22% of 100 random samples of the same number of putatively neutral SNPs were found in ancient tracts (Supplementary figure 22). The spread of these ancient coastal alleles by contemporary gene flow between coastal populations, including between the east and west sides of the North Atlantic is very unlikely given the site fidelity and low dispersal of coastal bottlenose dolphins (27, 36). These ancient alleles could have been introgressed from a unsampled “ghost” population which diverged from the sampled populations a long time ago, so that the introgressed regions contain mutations which accumulated in the ghost population over time, likely close to the split time between *T. truncatus* and *T. aduncus*. However, we do not have further support for this hypothesis and it is difficult to explain how this could have happened in different oceanic basins.

Rather, a most parsimonious hypothesis given the prevalence of these SNPs as SGV in the pelagic populations, is that this adaptation likely occurred in different oceans by repeated selection on SGV, which persisted at low frequencies in the large pelagic population. There are precedents for such recurrent use of standing genetic variation in nature; ancient polymorphisms have enabled rapid repeated parallel ecotype formation in saltmarsh beetles (16) and in threespine sticklebacks (20, 54). In sticklebacks, freshwater adapted alleles have persisted as SGV in the large marine populations as a result of episodic recurrent gene flow from freshwater populations (the so-called Transporter Hypothesis) (20, 55).

We propose a similar mechanism in our study system, in which coastal-associated ancestry could have been retained at low frequency as SGV in the pelagic ecotype through episodic gene flow from coastal populations, possibly occurring during cyclical expansion and contraction of habitat during past climate shifts. This may allow new coastal ecotypes to rapidly and recurrently arise through selection acting upon ancient SGV maintained at low frequency in pelagic populations, when new coastal habitat becomes available, for example during interglacial periods. The TMRCA of the ancient tracts suggests that the coastal and pelagic allelic divergence occurred near to the divergence of the *T. truncatus* and *T. aduncus* lineages. Episodic admixture between ecotypes may be a recurrent process, which has happened throughout the evolution of common bottlenose dolphins and pre-dates the formation of the present-day coastal ecotypes (Supplementary table 4). We see this akin to the ‘sieving’ of balanced polymorphism during the speciation process proposed by Guerrero and Hahn 2017 (17). Altogether, our results contribute towards the emerging hypothesis that old polymorphisms may allow rapid ecotype formation when new ecological opportunities arise, and ultimately ecological speciation (19).

Our results together with previous studies on human populations represent rare examples of species with long generation time for which parallel evolution has been uncovered (7, 21). In humans, parallel adaptation was facilitated by similar stable lifestyles (e.g. life in high altitudes (7)), or same cultural revolutions (e.g. cattle domestication for lactase persistence (21)). Non-human examples of socially driven local adaptation are scarce, but killer whale ecotypes have likely evolved as a result of demographic history, ecological opportunity and cultural transmission (24). Bottlenose dolphins (*Tursiops* sp.) also exhibit complex behaviors, such as habitat specialisation or social learning of foraging techniques, that strongly influence their patterns of genetic variability (28, 31, 38), and we hypothesize that these also facilitate their ability to adapt to novel conditions.

Although further investigation is warranted as many complex traits may be polygenic (56), and it is difficult to prove causal relations between behavioral/ecological traits and genes under selection, we uncovered parallel selection in 45 genes including some related to behavioral and ecologically relevant functions (Supplementary table 7). We found genes related to cognitive abilities, learning and memory (RELN (57, 58), ADER3 (59)), neuronal activity regulation (INSYN2A), lipid metabolism (AGK, LPIN2 (60), KLB), muscle contraction (RYR1, myosin-3, myosin-13, CAMK2D), axe growth (FEZ1, FEZ2), heart functions (CAMK2D), tooth enamel development (MMP20 (61)), immunity (HLA class II histocompatibility antigen, DQ alpha 2 chain, SERINC5), oxidative stress (cytochrome b5 reductase 4) and hormone regulation (STAR).

Perhaps the most interesting genes are those involved in cognitive, learning and memory abilities (RELN and ADER3) (57–59). RELN encodes for the reelin protein, which has a role in the modulation of synaptic transmission in response to experience (57, 58). Bottlenose dolphins (*Tursiops* sp.) have a propensity for innovation, i.e. developing habitat specific foraging techniques, which are transmitted maternally or in social groups (38, 62, 63). Socially transmitted foraging behaviour and complex coastal environments harboring a mosaic of prey may require genetic adaptations for increased cognitive abilities. RELN has been found under positive selection in sea otters, which also exhibit maternally transmitted prey preferences and tool use (64).

Other ecologically relevant genes include those involved in lipid metabolism and storage (AGK, LPIN2, KLB (60)), which may be involved in adaptation to the differing diets documented in coastal, mainly involving large fish, and pelagic, primarily pelagic fish and squid, populations (31, 33). Physiological demands (65) and food resource availability are different in coastal and pelagic environments, possibly leading to different constraints on lipid storage and fat-mass body composition controlled by LPIN2 (60). Parallel adaptation in immunity related genes (HLA class II histocompatibility antigen, DQ alpha 2 chain, SERINC5), may highlight different pathogen load between coastal and offshore environments.

We observed that 113,530 SNPs were under divergent selection among coastal populations, potentially highlighting coastal population specific divergent selection. This is not surprising given the heterogeneity of the different coastal habitats across the bottlenose dolphins’ range. Coastal dolphins are smaller than their pelagic counterparts in the NWA, while the pattern is opposite in other regions (34). This may be the result of differing selective pressures such as temperature dependent morphology. Our findings corroborate other studies challenging the binary view of parallel *vs*. non-parallel evolution, and point towards a continuum (12–14). This holds even for the most emblematic example of parallel evolution, the threespine sticklebacks, where deviation from parallel adaptation may be the result of geographic distance, stochastic processes and adaptation to environmental variation within habitat types (13, 14).

## Conclusion

Our results show that selection acted upon ancient SGV fueling parallel adaptation of common bottlenose dolphins to coastal environments, which also involved divergent selection among different coastal habitats. Repeated bouts of selection on genetic variation may promote adaptation to coastal habitat via re-using linked variants with minimal pleiotropic effects, thereby facilitating their persistence at low frequency in source populations and enabling parallel evolution of derived populations at the range margins (66). Stable transmission of ecological specialisations and a propensity for innovation and learning possibly facilitated local adaptation. Our study contributes to the growing body of evidence that ancient polymorphisms are a major substrate for rapid ecological adaptive divergence (19) and can have a key role in local adaptation of long-lived organisms. Therefore, they could help species to cope with environmental changes driven by current global change.

## Material and methods

### Sample collection and laboratory procedures

Epidermal tissue samples were collected from 57 bottlenose dolphins (Supplementary text, Figure 1, Supplementary table 1).

### Laboratory procedures

DNA extraction protocol procedures are detailed in the Supplementary text. Library and whole genome re-sequencing was performed at Beijing Genomics Institute (BGI). Illumina libraries were built on 300-bp DNA fragments and sequenced on an Illumina HiSeq X Ten platform (Supplementary text).

### Data processing and filtering

The read trimming and mapping, and data filtering are described in detail in the Supplementary text. Sequencing reads were processed with Trimmomatic v. 0.32 (67) using default parameters and sequence reads shorter than 75 bp were discarded. The remaining filtered reads were first mapped to a modified version of a published common bottlenose dolphin mitochondrial genome (GenBank KF570351.1) (68). Reads that did not map to the mitochondrial genome were then mapped to the reference common bottlenose dolphin genome assembly (GenBank: GCA_001922835.1, NIST Tur_tru v1) using BWA mem (v. 0.7.15) with default options (69).

Picard-tools v. 2.1.0 (70) was used to add read groups, merge the bam files from each individual from the different lanes and remove duplicates reads. Then, indel realignment was performed using GATK v. 3.6.0 (71). We kept only the mapped reads with a mapping quality of at least 30 and removed repeated regions as identified using RepeatMasker (72), regions of excessive coverage and the sex chromosomes (see details in the Supplementary text).

### SNP calling using genotype likelihoods

We called SNPs taking genotype uncertainty into account by calculating genotype likelihoods in ANGSD v. 0.913 (73) and keeping SNPs with a minimum MAF of 0.05 and having data in a minimum of 75% of the individuals. In ANGSD, all analyses described below were run considering only SNPs with a phred quality and a mapping quality score of 30. We further filtered the dataset by excluding SNPs that showed significant deviation from Hardy-Weinberg Equilibrium (HWE) and an inbreeding coefficient (F) value <0 as this can also be the result of paralogues or other mapping artefacts.

### Linkage Disequilibrium (LD) pruning and population structure

We excluded one individual from the population structure analyses that were not based on population allele frequencies as it had a coverage much lower than the others (sample 7Tt182 from the NWAp population). We used NgsLD (74) to obtain a set of unlinked SNPs (Supplementary text). Population structure analyses, admixture analysis in NGSAdmix (40) and PCAs in PCAngsd (39) were run using a set of 798,572 unlinked SNPs. NGSAdmix was run 10 times for each *K* value between 2 and 8, using a tolerance for convergence of 1e^−10^ and a minimum likelihood ratio value of 1e^−6^. Consistency between runs was checked and the runs with the highest likelihood were plotted. The highest level of structure was identified using the Evanno method (75).

### Ancestral state reconstruction

We describe how we reconstructed the ancestral state of alleles in the Supplementary text.

### Genotype calling

We called variants (i.e. generation of a vcf file) using samtools v. 1.2 mpileup and bcftools multiallelic and rare-variant calling, option –m (76, 77) on the filtered bam files. Variable sites with a minimum mapping quality of 30, a phred score quality of 30 and genotype quality of 20 were retained in vcftools (78). We kept SNPs with a minimum MAF of 0.05 and having genotype data in a minimum of five individuals in each of the seven populations. The vcf file was also filtered for monomorphic and non-biallelic sites, totaling 2,003,833 SNPs. A vcf file was used as an input for the analyses described below apart from the SFS and diversity estimates, which were estimated using genotype likelihoods in ANGSD.

### Demographic history

We computed demographic history, that is changes in effective population sizes (*N*_e_) through time and ecotype splits within a region and splits of the different pelagic ecotypes, using the program SMC++ (42). Details of the analysis procedure, run on autosome scaffolds which were more than 10 Mbp, and on a vcf file not filtered for any MAF, are provided in the Supplementary text. Briefly, the repeated regions and excessive coverage regions were included as a mask file so that they were not misidentified as very long runs of homozygosity. The analysis was run both using all regions and taking out all the regions under selection, as identified with Flink (see below). Regions under selection were defined as 50kb around each outlier SNPs. Regions under selection were included in the mask file when they were taken out from the dataset. Population size histories and splits times between ecotypes in each region and between the pelagic populations were estimated using the default settings, a generation time of 21.1 years for the species (79) and two different mutation rates. Mutation rates were i) 9.10e-10 substitutions per site per year that is 1.92e-8 substitution per nucleotide per generation (80) and ii) 1.21e-9 substitution rate per site per year (81) that is 2.56e-8 substitution per nucleotide per generation. Results were plotted in R v. 3.6.1 (82) (Supplementary text).

### Diversity and population structure statistics

We estimated the unfolded site frequency spectrum (SFS), the 2D-SFS, nucleotide diversity, Watterson’s Theta and Tajima’s D for each population using ANGSD (see details in the Supplementary text). We calculated nucleotide diversity and Watterson’s Theta for each site and then we estimated both the latter and Tajima’s D using a sliding-window size of 50 kb and a step size of 10 kb. We estimated pairwise weighted *F*_ST_ using vcftools (78) using a sliding-window size of 50 kb and a step size of 10 kb.

### Admixture analyses

We reconstructed the relationships of the coastal and pelagic ecotypes from the different regions as a Maximum Likelihood bifurcating tree using TreeMix version 1.13 (44) (Supplementary text). We ran TreeMix using one individual *Tursiops aduncus* (SRX2653496/SRR5357656 (83)) as a root. Reads of this *T. aduncus* individual were mapped to the common bottlenose dolphin reference genome assembly as described above and processed as described earlier for our data. As we had one individual for this species, we used one randomly chosen individual from each of our populations. We ran TreeMix with three different sets of individuals to check consistency of the results when including different individuals. We first ran TreeMix ten times for each value of m (migration events) ranging from 0 to 10 (-noss -global -k 1000). We estimated the optimal number of migration events to 1 using the optM R package (https://cran.r-project.org/web/packages/OptM/index.html). We then ran TreeMix 100 times for 0 (as null model) and 1 migration event and obtained a consensus tree and bootstrap values using the BITE R package (84). The residual covariance matrix was estimated for each m value and the consensus tree using TreeMix.

We then estimated *F*_4_-statistics to test whether geographic pairs of pelagic and coastal ecotypes had evolved independently (45, 46). The *F*_4_-statistics can be used to test whether a given tree describes accurately the relationships among four test populations and to detect admixture events (see Supplementary text for details). *F*_4_-statistics were computed for each possible combination of population using the fourpop function in TreeMix version 1.13 (44). We accounted for linkage disequilibrium by jackknifing in windows of 1,000 SNPs. This block jackknife was used to obtain a Standard Error (SE) on the estimate of the *F*_4_-statistics and test for significance using a *Z*-score.

### Ancient ancestry analyses

Ancient tracts introgressed into the coastal ecotype from a divergent lineage after splitting from the pelagic source population, or differentially sorted from structure in an ancestral population will contain clusters of private alleles, the density of which will depend upon the divergence time of the introgressing and receiving lineages (49, 85) (Supplementary figure 13, Supplementary text). We therefore set out to screen for genomic tracts of consecutive or clustered private (i.e. relative to the allopatric pelagic individuals) alleles in each of the individuals from the coastal ecotype. To ensure the results are comparable despite variation between samples in coverage at some sites, we randomly sampled a single allele at each site from each diploid modern genome in all scaffolds longer than 1Mb using ANGSD. For the outgroup we used all variants found in a dataset consisting of all non-allopatric pelagic samples (Supplementary figure 13). We then used a Hidden Markov Model (HMM) to classify 1 kb windows into ‘non-ancient’ and ‘ancient’ states based on the density of private alleles (47), see details in Supplementary text. We considered windows inferred as ancient as those with posterior probabilities of P>0.8 (47, 49). We also estimated the mean TMRCA of the ancient and non-ancient (ingroup) windows with the corresponding segments in the outgroup dataset.

### Patterns of structuration across the genome

We used localPCA (52) to describe the three major patterns of relatedness (“corners”) among populations on four PCs for each scaffold longer than 10 Mbp using the default options (two PCs and two MDS coordinates), the R codes available on github and bins of 100 SNPs. We plotted the pairwise plots of the first four PCs for each of the three corners.

### Selection analyses

We used Flink (41) to test for selection to coastal *vs*. pelagic habitat. Flink is an extension of Bayescan (53), respectively describing selection and drift, which takes linkage among loci into account. Specifically, it applies a hidden Markov Model to identify the effect of selection at linked markers using correlation in the loci specific elements along the genome. Flink was run grouping the populations into two groups: coastal and pelagic. Each group was composed by the three populations from each region. Scaffolds were grouped into super-scaffolds, so that each contains at least 50,000 SNPs. In Flink, the function estimate was run, and parameters settings are described in the Supplementary text. The number of iterations was set to 500,000, the burnin to 300,000 and the thinning to 100. We considered a locus under selection when it is within the 1% FDR threshold.

To get further insights into the results obtained by Flink, we plotted the raw genotypes of all the SNPs, SNPs under homogenising selection in the coastal populations, SNPs under divergent selection between ecotypes, and SNPs under both homogenising selection in the coastal populations and divergent between ecotypes (defined as the SNPs under parallel selection) in R (see details in the Supplementary text). We also plotted a neighbor-joining distance tree and a PCA for the SNPs under each type of selection. To determine the origin of the SNPs under selection, we defined how many were also polymorphic in the pelagic populations, and compared the 2DSFS between all pairs of populations, estimated in ANGSD (see details in Supplementary text). Then, we defined how many SNPs under the different types of selection were found in ancient tracts. We compared the results with 100 random samples of the same number of putatively neutral SNPs found in ancient tracts.

We identified the genes associated with the SNPs under parallel selection using the reference genome annotation file. We describe how we determined the putative functions of the genes under selection in the Supplementary text.

## Acknowledgements

We thank everyone who was involved in sample collection. We thank Joseph Ward and Peter Thorpe from the St Andrews Bioinformatics Unit for help with software installations and troubleshooting. Bioinformatics and Computational Biology analyses were supported by the University of St Andrews Bioinformatics Unit which is funded by a Wellcome Trust ISSF award [grant 105621/Z/14/Z].

## Ethics

Samples were collected under permits MMPA Permit 779-1633 and MMPA Permit 779-1339 for the NWA and NMFS 14097 for the NEP. They were shipped from the Southwest Fisheries Science Center, NOAA Fisheries, USA to the University of St Andrews, UK under CITES institutional permits, US057 and GB035, and the US Fish and Wildlife Service permit 16US690343/9 and to BGI in Hong Kong, China under CITES export permit 547016/01.

## Author contributions

ML initiated the study, ML, MCF, ADF, OEG conceived the analyses. FA, SB, AB, JOB, KR and PER collected the samples. KR, PER and MCF did the DNA extractions. ML, MG, MN, RF, BSB, MCF, ADF and OEG ran the analyses, and ADF, OEG and MCF provided supervision. ML wrote the manuscript with input from all co-authors.

