## Supplementary material for "Selection on ancestral genetic variation fuels parallel ecotype formation in bottlenose dolphins"

Supplementary material – text, figures and tables

Louis M., Galimberti M., Archer F., Berrow S., Brownlow A., Fallon R., Nykänen M., O'Brien J., Roberston K. M., Rosel P. E., Simon-Bouhet B., Wegmann D., Fontaine M.C.*, Foote A.D.*, Gaggiotti O.E.*

* these authors contributed equally

### Extended results

#### Population structure

PCA (1) and NGSAdmix (2) analyses run on a set of 798,572 unlinked SNPs indicated that the three pelagic populations were more closely related to each other than the three coastal populations which were much more differentiated from each other (Figure 1, Supplementary figures 1-3). The NGSAdmix analysis showed a strong increase in logP between $K$ = 3 and 4, which was also the greatest change in mean log likelihood (3) and almost no increase after $K$ = 6 (Supplementary figures 1a-b). With $K$ = 4, all pelagic populations were grouped together while each coastal one represented a distinct cluster. With $K$ = 6, each population represented a distinct cluster, but the North-East Atlantic (NEA) pelagic population exhibited some admixture with the two other pelagic populations (Supplementary figure 1c). Atlantic populations separated out from Pacific populations along PC 1 (10.2% of the variance explained, Supplementary figures 2-3), with the greatest difference between North-East Pacific coastal (NEPc) and North-West Atlantic coastal (NWAc). Coastal populations segregated from pelagic populations along PC 2 (8.45% of the variance explained), with the greatest difference along this axis being between the coastal and pelagic ecotypes in the North-West Atlantic (NWAc *vs.* NWAp). The location of the samples within the PCA plot can be interpreted in terms of the mean pairwise coalescent time between each pair of samples (4), so that samples with greater covariance of alleles than the mean covariance among samples cluster together. Samples from the same population clustered together. These results thus suggest greater covariance of alleles among populations within the same ocean basin, and greater covariance of alleles among the pelagic populations than among coastal populations. The location of the coastal samples on the PCA suggests several founding events of coastal populations. Shared variation between the two pelagic populations in the Atlantic indicated that the two Atlantic coastal populations were possibly independently derived from the same ancestral Atlantic population (Supplementary figures 1c, 3 and table 2).

**SMC++**

Coalescent-based estimates of effective population size (5) on putatively neutral regions found a concurrent decrease in $N_e$ from 400,000 to 150,000 yBP (years Before Present), followed by a steady increase at the onset of the last glacial period between 100-150,000 yBP until 50,000 yBP (Figure 2a, Supplementary figures 4, 5a-b). It is important to note that these plots reflect changes in coalescence rates through time, which could also indicate changes in population structure rather than changes in the number of individuals (6). From approximately 50,000 yBP pelagic and coastal populations showed contrasting demographic histories. Pelagic populations in the Atlantic had relatively constant $N_e$ after 50,000 yBP and showed similar demographic trajectories, while the NEP pelagic population kept increasing in size until around 4,000 yBP. The NEA and NEP coastal populations declined in $N_e$ from around 50,000 yBP, followed by post-glacial expansion. The NWA coastal population showed a disjoint demographic history with the NWA pelagic population at the onset of the LGM. This is shortly followed by the estimated divergence time between the two ecotypes in the NWA at around 80,000 yBP (Figure 2c, Supplementary figure 11a-b). Divergence time between the two ecotypes in the Pacific region was estimated to be around 50,000 yBP. The divergence of the NEA coastal and pelagic ecotypes occurred after the end of the Last Glacial Maximum. In contrast, the divergence time between the NEA and NWA coastal populations was estimated around 50,000-70,000 yBP (Supplementary figure 12a-b).

**TreeMix**

Demographic history and potential admixture events were explored using TreeMix (7), in which shared co-variance of drift of allele frequencies suggest admixture events (Supplementary figures 8-10). The results showed that a simple bifurcating tree without migration or admixture provided a poor fit to the data (Supplementary figure 8). Inspecting the residual covariance in allele frequencies between pairs of populations (Supplementary figure 9) showed that the model was unable to explain properly allele covariance between the North-West Atlantic coastal population and an outgroup species, the strictly coastal Indo-Pacific bottlenose dolphin, *Tursiops aduncus*. These strongly positive residuals for the NWAc and *T. aduncus,* and to a lesser extent between NEAp and NEPp, NEAp and NWAp, NWAp and NEPp, NEAc and NEPc and, NEAc and NEPp in the tree without migration indicated they are more closely related to each other than in the best-fit tree and are candidates for admixture. The best number of migration edges was estimated to m=1, using an ad hoc analysis of the second order rate of change in the log-likelihood (Evanno) method (Supplementary figure 8). Adding a single migration event improved the model fit to the data, while additional migration edges did not provide further improvement (Supplementary figure 8). Ninety-nine percent of the variance was explained using m=1, and variance only increases marginally at m>1. However, the addition of this migration edge resulted in incongruence among *T. truncatus* populations, indicating the admixture graph was still not a good fit for the data.

The maximum likelihood population tree estimated with TreeMix (7) with or without migration clearly supported the independent split of the Pacific and Atlantic coastal populations (Supplementary figures 9 and 10). In contrast, the two coastal populations in the NWA and NEA were closely related. The TreeMix results thus suggest the contribution of an outgroup to the genetic composition of the NWA coastal population and admixture among our studied populations.

We found more segregating sites in the NWAc population in comparison to other populations (Supplementary figures 16-17b, Supplementary table 3), indicating private ancestry in this population. If the source of this variation is admixture from an independent, unsampled, lineage of *T. truncatus*, it could be represented as a migration back to *T. aduncus* from NWAc by Treemix. D-statistics (8–10) did not support introgression between NWAc and *T. aduncus* and indicated that the NWAp population is closer to *T. aduncus* than NWAc (Supplementary table 6), possibly due to extended period of drift in the NWAc population. The position of the NEP populations having slightly less drift from the basal node than the NA populations could be due to small amounts of gene flow between *T. aduncus* and the NEP samples, both species occurring in the western Pacific. D-statistics do not support preferential introgression of *T. aduncus* with the NEPc or NEPp (Supplementary table 6). Alternatively, shared ancestral polymorphisms that differentially segregate in coastal *T. truncatus* and *T. aduncus* lineages may explain the migration edge from the NWAc to *T. aduncus*, see Discussion.

Overall, the TreeMix results show a lack of clear topology among the *T. truncatus* populations. These results and our other admixture test results reinforce the idea that these highly mobile marine populations do not conform to a simple bi-furcating tree model (11, 12). We therefore chose to apply mainly population genomic methods, which investigate variation in allele frequencies and account for complex ancestry, rather than phylogenetic substitution-based inferences that assume a simple bifurcating branching process.

In addition, the TreeMix model has a number of assumptions that should be taken into account when interpreting the results (7). The model assumes a mainly tree-like history of populations and considers that migration occurred at a single point in time. In this study, the covariance in allele frequencies between populations may be a consequence of low levels of on-going gene flow between populations at equilibrium rather than discrete migration events.

**Ghost ancestry**

The TMRCA between the ancient tracts in coastal dolphins and the corresponding genomic regions in the outgroup had a Poisson distributed mean between 40 and >100 thousand generations (~845-2,257 KYA, assuming a generation time of 21.1 years (13), Supplementary table 4) depending upon the mutation rate assumed. $T_{\text{Ancient}}$ was the oldest for the NWAc ecotype, which also had approximately double the proportion of inferred introgressed ancestry within the genome (>20 Mb) at a posterior probability of P>0.8, compared to the NEAc and NEPc populations (Supplementary table 3). This suggests a greater action of recombination upon introgressed ancestry in the NEAc and NEPc populations.

**Flink**

Overall, selective pressures were the most pronounced among coastal populations (Figure 4a). The coastal ecotype also exhibited the largest proportion of loci under homogenising selection. On the other hand, no loci under homogenising selection were detected between coastal and pelagic dolphins. The proportion of loci under homogenising selection in pelagic dolphins was significantly lower than that observed in coastal dolphins. In terms of divergent selection, its incidence in coastal populations was similar to that observed between coastal and pelagic populations, with the pelagic population exhibiting the lowest incidence. As opposed to coastal dolphins, pelagic populations exhibited a similar proportion of neutral loci as that observed between coastal and pelagic ecotypes.

Almost all SNPs under homogenising selection in the coastal populations were also polymorphic in the pelagic populations (96% of in all the pelagic, 83% in the NEAp, 82% in the NEPp, 78% in the NWAp), supporting natural selection having acted mostly upon Standing Genetic Variation (SGV). Plots of the 2DSFS also show that there was more shared variation between a coastal and pelagic population within a region than when looking at all the sites (Supplementary figure 18). They also indicate more shared variation between the coastal populations (Supplementary figure 18), although there were still a lot of private alleles. Genotypes of the SNPs under homogenising selection in the coastal populations were more similar between ecotypes (Supplementary figure 19a) than those of all the SNPs (Supplementary Figure 16-17a). In the PCA ran on the SNPs under homogenising selection in the coastal populations, the pelagic populations clustered together, and coastal populations were differentiated from them and from each other (Supplementary figure 19b). The first axis separated the Pacific and the Atlantic. In the unrooted NJ tree (Supplementary figure 19c), generated from a distance matrix computed in the adegenet package in R, the two Atlantic populations clustered together, while the NEPc population clustered independently.

Most of the SNPs under divergent selection between the ecotypes were also polymorphic in the pelagic populations (72% of in all the pelagic, 56% in the NEAp, 55% in the NEPp, 51% in the NWAp), also supporting that selection has acted mostly upon SGV. Plots of the 2DSFS show that there was less shared variation between a coastal and a pelagic population within a region, than in all sites (Supplementary figure 18). In contrast, there was more shared variation among coastal populations than in all sites (Supplementary figure 18). Genotypes of the SNPs under divergent selection between ecotypes were more dissimilar between the coastal and pelagic populations (Supplementary figure 20a) than those of all SNPs (Supplementary Figure 16-17a). There was more fixed variation overall in the coastal populations, apart in the coastal population of the NEA, where there were many heterozygotes. The genotypes of the NWAc population standed out as mainly fixed for the reference genome (which is a coastal individual). In the PCA ran on those SNPs, the pelagic populations clustered together with the NEPc, and the two coastal populations in the Atlantic were differentiated from them and from each other (Supplementary figure 20b). In the unrooted NJ tree, the dolphins clustered by ecotype, all coastal populations clustered together, and likewise for the pelagic populations (Supplementary figure 20c).

Most of the SNPs under both homogenising selection among coastal populations and under divergent selection between the ecotypes (i.e. under parallel selection) were also polymorphic in the pelagic populations (81% of in all the pelagic, 61% in the NEAp, 65% in the NEPp, 58% in the NWAp), again supporting selection acting mainly upon standing genetic variation. The plot of the genotypes clearly shows different patterns in the coastal and the pelagic populations (Figure 5a). Genotypes were more heterozygotes in the coastal populations. In the PCA ran on those SNPs, the coastal populations were separated from the pelagic populations on the first axis with individuals from the same coastal population clustering together and apart from the other coastal populations on the second axis (Figure 5b). In the unrooted NJ tree, the dolphins clustered by ecotype; all coastal populations clustered together, and so did the pelagic populations (Figure 5c).

Thirty-three percent of the SNPs identified in the region as under homogenising selection were found in ancient tracts, while 61 and 66% of the SNPs under divergent selection between ecotypes, and under both types of selection were respectively found in ancient tracts (Supplementary figure 22). In contrast, only an average of 20%, 20% and 22% of putatively neutral SNPs were found in ancient tracts in 100 random samples of the same number of SNPs in homogenising, divergent and parallel selection, respectively (Supplementary figure 22).

### Extended material and methods

#### Sample collection and laboratory procedures

Epidermal tissue samples were collected from 57 bottlenose dolphins (Figure 1, Supplementary table 1). Samples from the NEA were stranded animals found dead on the beach and assigned to the coastal and pelagic ecotypes in a previous study using microsatellites and a portion of the mitochondrial control region data (14). Coastal dolphins included five previously photo-identified bottlenose dolphins on the east coast of Scotland. Samples from the NWA and the NEP were collected as per Rosel *et al.* 2009 (15) and Lowther-Thieleking *et al.* 2015 (16) respectively. Samples were previously assigned to either coastal or pelagic ecotypes (14). Samples were stored at −80°C with no preservative or at −20°C fixed in a salt-saturated 20% DMSO solution or ethanol or with no preservative.

DNA was extracted from epidermal tissue using a Qiagen DNeasy kit following the manufacturer's protocol for the NEA samples. For the NWA samples, DNA was extracted using standard proteinase K digestion and organic extraction as described in Rosel and Block 1996 (17). For the NEP samples, DNA was extracted using a sodium chloride protein precipitation (18).

Illumina libraries were built on 300-bp DNA fragments and were pooled equimolarly by geographical regions. The geographic region pools were sequenced across six lanes of an Illumina HiSeq X Ten platform for the NEA and NEP (20 DNA samples each) and five lanes for the NWA (17 DNA samples) using paired-end 150-bp chemistry.

#### Read trimming and mapping

Demultiplexing was performed by BGI. Sequencing reads were processed with Trimmomatic v. 0.32 (19) to trim residual adapter sequence contamination using the default options for the seed mismatches: 2, the palindrome clip threshold: 30 and the simple clip threshold: 15 and low quality bases. We removed low quality bases (a phred score of less than 5) from the beginning and the end of the reads. In addition, we also performed a sliding-window trimming, cutting bases when the average quality within a window of 4 bp fell below a phred score of 15 (default parameter). Sequence reads that were less than 75 bp were discarded.

The remaining filtered reads were first mapped to a bottlenose dolphin mitochondrial genome (Genbank gi_557468684_gb_KF570351.1_.fasta) (20) as per Morin et al. 2015 (21). Reads that did not map to the mitochondrial genome were then extracted from the bam file and converted into a fastq file using samtools v. 1.2 (22, 23) and picard-tools v. 2.1.0 (24). These reads were then mapped to the reference bottlenose dolphin genome assembly (Genbank: GCA_001922835.1, NIST Tur_tru v1) using BWA mem (v. 0.7.15) with default options (25).

**Data filtering**

We checked and confirmed the quality of our data using FASTQC (26) after trimming, mapping and filtering. Picard-tools v. 2.1.0 (24) was used to add read groups and merge the bam files from each individual from the different lanes and remove duplicates reads. The optical duplicate pixel distance was set to 2,500 as recommended by the Broad Institute to better estimate library complexity on data generated using the Illumina HiSeq XTen platform. Then, indel realignment was performed using GATK v. 3.6.0 (27, 28). Samtools was used to keep only the mapped reads with a mapping quality of at least 30.

Repeat regions from the cetartiodactyla group were identified in the bottlenose dolphin reference genome using RepeatMasker (29) and saved in a bed file. Only interspersed repeats were masked and STRs, small RNAs and low complexity regions were retained. The repeat regions were removed from the bam files using bedtools v. 2.25.0 (30) and samtools v. 1.2. We also removed regions of excessive coverage as the high coverage of these regions can potentially be the result of unmasked repeated regions, in nuclear mitochondrial DNA (NUMTs), or some other mapping artifact (e.g. paralogous loci). Coverage was then estimated for each genome using the doDepth function ANGSD v. 0.913 (31). However, due to the size of the dataset, global coverage was estimated by randomly sampling three individuals per ecotype (i.e. for a total of 18 individuals). Regions that were higher than twice the mean coverage (>346x) were considered of excessive coverage. These regions were detected using the CALLABLELOCI tool in GATK. Then they were removed from the bam files using bedtools and samtools as above. Mean coverage was again estimated using the doDepth function in ANGSD.

To identify the scaffolds corresponding to the autosomes and the X chromosomes, we randomly sampled five males and five females and estimated mean coverage for males and females for all scaffolds that were longer than 1 Mbp (i.e. 98.3% of the genome). When the ratio of the mean coverage for the females on the mean coverage for the males was around 1 (mean=1.06, min=1.04, max=1.19) it was considered the scaffold corresponded to an autosome and when the ratio was around 2 (mean=2.05, min=1.96, max=2.07), the scaffold was considered as belonging to the X chromosomes. Twelve scaffolds > 1Mbp were identified as corresponding to the X chromosomes and 116 scaffolds to the autosomes. The 12 scaffolds belonging to the X-chromosomes were removed using bedtools and samtools.
When the sex was unknown, it was identified by estimating coverage for the 12 X-chromosome scaffolds and 12 scaffolds from the autosomes.

ANGSD was used to identify SNPs that show significant deviation from HWE and a F value <0. The latter sites were removed using bedtools and samtools as above. Sites with a F>0 were kept as this may indicate Wahlund effect.

**Linkage Disequilibrium (LD) pruning**

NgsLD (32) was used to obtain a set of unlinked SNPs. It was first run with a maximum distance of 1,000 Kb for SNPs to be possibly in LD, randomly subsampling 5% of the data and LD decay was inspected using R. As LD decay was decreasing rapidly, NgsLD was then re-run using a maximum distance of 100 kb. LD decay was inspected and showed LD was negligible after a distance of 20 kb. Then, a set of unlinked sites was produced considering that SNPs are in LD until 20 kb and using a minimum weight of 0.5. Population structure analyses were run on the set of unlinked SNPs.

**Ancestral state reconstruction**

The ancestral state of the alleles was reconstructed by creating a consensus sequence using two whole genomes of the killer whales (*Orcinus orca*), the sperm whale (*Physeter* *macrocephalus*) and the finless porpoise (*Neophocaena phocaenoides*). Short read data from two killer whales, a sperm whale and a finless porpoise (SRR574982/SRX188934, SRR1162264/SRX447351, SRR1031998/SRX378812, SRR940959/SRX326372 (33–36)) were additionally accessed from the National Center for Biotechnology Information Sequence Read Archive database and mapped to the common bottlenose dolphin reference genome assembly as described above for the modern bottlenose dolphin samples. Slight changes included: i) setting the phred score to 20 instead of 15 for the sliding window in Trimmomatic, as base quality of the raw data were lower than for the bottlenose dolphin data and ii) setting the minimum mapping quality to 20 as mapping in against another species. Coverage was estimated for each genome using the doDepth function in ANGSD and they were then subsampled to a coverage of 5x using samtools. The four genomes were merged using samtools and then the consensus sequence was inferred by selecting the most common base using the doFasta 2 option together with doCounts 1 in ANGSD.

**SMC++**

Demographic history, that is changes in effective population sizes ($N_e$) through time and ecotype splits within a region and splits of the different pelagic ecotypes, were computed using the program SMC++ (5), i.e. Sequential Markov Coalescent + plenty of unlabelled samples. The method incorporates both the Site Frequency Spectrum method and the Linkage Disequilibrium information and recombination rate in a coalescent Hidden Markov Model, HMM (similar to PSMC, Pairwise Sequentially Markovian Coalescent (37), and MSMC (38), Multiple Sequentially Markovian Coalescent). PSMC uses the distribution of heterozygous sites throughout the genome where the heterozygosity information is emitted as binary. In addition, SMC++ emits the allele frequency of an extra n-2 haplotypes. The latter is based on the SFS conditioned on the TMRCA of a single distinguished individual. It can include several individuals per population while PSMC analyses are only based on one individual. In contrast with MSMC, phasing the data is not required.

Then, only the autosome scaffolds, which were more than 10 Mbp were included in the analyses, and no MAF filter was applied on the vcf file. The vcf file was converted to SMCpp format using the vcf2smc function for each retained scaffold. The repeated regions and excessive coverage region were included as a mask file so that they were not misidentified as very long runs of homozygosity which could impact the population trajectories and create false recent sudden decreases in $N_e$ in recent times. The analysis was run both using all regions and taking out all the regions under selection, as identified with Flink (see below). Regions under selection were defined as 50kb around each outlier SNPs (thus 25 kb each side). Regions under selection were included in the mask file when they were taken out from the dataset.

We fixed the distinguished individual to i) a particular individual, or ii) made it vary over two individuals and iii) made it vary over three individuals. Varying the distinguished individual over different individuals has the advantage of incorporating genealogical information from additional individuals into the analysis, which may lead to improved estimates.

Population size histories were estimated using the *estimate* option in SMC++ using the default settings for the estimate function, a generation time of 21.1 years for the species (13) and two different mutations rate. Mutation rates were i) 9.10e-10 substitutions per site per year that is 1.92e-8 substitution per nucleotide per generation (39) and ii) 1.21e-9 substitution rate per site per year (35) that is 2.56e-8 substitution per nucleotide per generation.

Population split estimations first involves estimating population histories using the estimate option, i.e. the marginal estimates. Then, datasets containing the joint frequency spectrum for both populations were computed using the vcf2smc function. Lastly, the *split* function was used to refine the marginal estimates into an estimate of the joint demography and divergence times were estimated between ecotypes in each region and between the pelagic populations. Results were plotted in R v. 3.6.1 (40) with packages ggplot2 (41), scales (42) and RcolorBrewer (43)).

**Diversity estimates**

Nucleotide diversity, Theta Watterson and Tajima D were estimated for each population using ANGSD (31, 44). First the unfolded site frequency spectrum (SFS) was computed for each population in ANGSD using a two step procedure (44) for sites with data in all individuals and including the ancestral state as defined earlier. First, the dosaf 1 function was used to calculate the site allele frequency spectrum likelihood (saf) based on individual genotype likelihoods assuming HWE. Then, the realSFS function was used to optimize the saf and estimate the SFS. Nucleotide diversity and Theta Watterson were calculated for each site and then both the latter and Tajima D were estimated from the SFS using a sliding-window size of 50 kb and a step size of 10 kb. To compute the 2D-SFS, the realSFS function was run on the saf files from each pair of populations.

**Admixture analyses**

We reconstructed the relationships among coastal and pelagic bottlenose dolphins in the NEA, NWA and NEP using admixture and 'treeness' tests. In contrast to STRUCTURE (45), which is suitable to detect population substructure but does not provide any test for admixture (i.e. the structure detected can be the result of multiple population histories), TreeMix and $F_4$-statistics explicitly test for admixture and can also inform on the directionality of gene flow (10).

TreeMix estimates a bifurcating ML tree based on genome-wide population allele frequency data and uses a Gaussian approximation to estimate genetic drift among populations. The relationships among populations are represented by the branches of the tree based on the majority of alleles. Migration edges are fitted between populations that are a poor fit to the tree model from the covariance matrix of allele frequencies. The direction of the migration events is inferred from asymmetries in the covariance matrix of allele frequencies relative to an ancestral population.

The $F_4$-statistics can be used to test whether a given tree describes accurately the relationships among four test populations and to detect admixture events even if they occurred hundreds of generations ago (46). It is robust to incomplete lineage sorting and changes in effective population size (46). The $F_4$-statistics quantifies drift (i.e. changes of allele frequencies) between pairs of populations in a tree (10, 46). The relationships between four populations can be described by three possible unrooted trees. For example, the relationships between populations A, B, C, D could be represented by three trees (A,B;C,D), (A,C;B,D) and (A,D;B,C). When the topology is correct, the difference in allele frequencies (i.e. the drift that has accumulated) between the two populations in each clade should be uncorrelated between clades (see details in Supplementary text). The $F_4$-statistics $F_4$(A,B;C,D) would thus not differ significantly from 0. For incorrect topologies, correlated drift would lead to significantly positive or negative correlation values. To test whether each geographic pair of pelagic and coastal ecotypes had evolved independently, we estimated $F_4$(pelagic$_x$, coastal$_x$; pelagic$_y$, coastal$_y$), and to test for a shared colonisation history of both the coastal and pelagic ecotype within a geographic region we estimated $F_4$(pelagic$_x$, pelagic$_y$; coastal$_x$, coastal$_y$).

D-statistics were estimated using ANGSD to assess the relationships of the NEP and NWA ecotypes to *Tursiops aduncus*. We tested these populations as *T. aduncus* is found in the Pacific, and the NWAc shows signatures of older ancestry, and we wanted to test whether this could represent ancient admixture with *T. aduncus.* D-statistics were calculated using one individual per population. The D-statistic describes an excess of shared derived alleles between taxa which could be the result of introgression or ancestral population structure. It thus allows to detect departure from 'tree-ness' of a given topology (8–10). We used the D-statistics to evaluate if the data are consistent with the null hypothesis that the tree (((H1,H2),H3),*Orca*) is correct and that there has been no past gene flow between *T. aduncus* and neither ecotype in a region. We used the killer whale (*Orca*) as the outgroup, and mapping is described above, as for the ancestral state.

The definition of the D-statistics used here is the one of Durand *et al.* 2011 (9).

D=(nABBA-nBABA)/(nABBA+nBABA) where nABBA is the number of sites where only H2 and H3 share a derived allele (ABBA sites) and nBABA is the number of sites where only H1 and H3 share a derived allele (BABA sites). Haploid sequences were generated for H1, H2 and H3 to avoid bias due to differences in coverage, by selecting a random base at each position. Under the null hypothesis that the given topology is the true topology, we expect an equal number of ABBA and BABA sites and thus D=0. A statistic differing significantly from 0 indicates either past gene flow between one population within the in-group and H3, or that the tree is incorrect. The significance from deviation from 0 was assessed using a *Z*-score based on blocked jackknife estimates of the standard deviation of the D-statistics (block size was 5Mb which should be higher than the LD in the populations). This *Z*-score relies on the assumption that the D-statistics, under the null hypothesis, is normally distributed with mean

0 and a standard deviation equal to a standard deviation estimate computed using the "delete-m jackknife for unequal m" method described in Busing et al. 1999 (47).

**Ancient ancestry analyses**

Ancient tracts with a distinctly older TMRCA than the genome-wide average, may result from introgression from an outgroup (48) or balanced polymorphism (Supplementary figure 13). Such tracts can be identified even without a reference genome of the introgression source. Private alleles resulting from *de novo* mutation along the branch to the coastal populations should be approximately randomly distributed across the genome. However, tracts introgressed into the coastal ecotype from a divergent lineage after splitting from the pelagic source population, or differentially sorted from structure in an ancestral population will contain clusters of private alleles, the density of which will depend upon the divergence time of the introgressing and receiving lineages (49, 50).

We therefore set out to screen for genomic tracts of consecutive or clustered private (i.e. relative to the allopatric pelagic individuals) alleles in each of the individuals from the coastal ecotype. To ensure the results are comparable despite variation between samples in coverage at some sites, we randomly sampled a single allele at each site from each diploid modern genome in all scaffolds longer than 1Mb. For the outgroup we used all variants found in a dataset consisting of all non-allopatric pelagic samples (Supplementary figure 13). We then used a Hidden Markov Model (HMM) to classify 1 kb windows into 'non-ancient' and 'ancient' states based on the density of private alleles (48). The background mutation rate was estimated in windows of 100 kb, using the variant density in all individuals. We then weighted each 1 kb window by the proportion of sites not masked by our RepeatMasker and CallableLoci bed files. The HMM was trained using a set of starting parameters based on those used for humans (48). We trained the model across five independent runs, varying the starting parameters each time to ensure consistency of the final parameter input. Posterior decoding then determines whether consecutive 1 kb windows change or retain state ('ancient' or 'non-ancient') dependent upon the posterior probability.

Considering windows inferred as ancient with posterior probabilities of >0.8 (48, 50), we identified > 1,000 ancient tracts totalling > 10Mb in each coastal genome tested (Supplementary table 3). The emission probabilities of the HMM are modelled as Poisson distributions with means of $\lambda_{Ancient} = \mu \cdot L \cdot T_{Ancient}$ for the introgressed state and $\lambda_{Ingroup} = \mu \cdot L \cdot T_{Ingroup}$ for the non-ancient (or ingroup) state (48), where $L$ is the window size (1000 bp) and $\mu$ is the mutation rate (1.92e-8 and 2.56e-8 substitution per nucleotide per generation (35, 39)). This allows us to estimate the mean TMRCA of the ancient and ingroup windows with the corresponding segments in the outgroup dataset.

**Flink**

In Flink (51), the function estimate was run with parameters A_max (maximum coefficient of selection) set to 4.0, B (coefficient of drift of a group) to a mean of -2.0 and standard deviation of 1.8, lnK (logarithm of positive scaling parameter) of a minimum of -10.0 and maximum of -0.1, lnMu (Probability involved in the generating matrix to go to a different state for the higher hierarchy) to -4.0, 0.0; lnNu (Probability involved in the generating matrix to go to a selection state from the neutral state for the higher hierar ) to -5.0, 0.0 - apart for scaffold groups 22 and 23 where it was reduced to -4.0 due to convergence issues, lnMu_g (Probability involved in the generating matrix to go to a different state at the group level) to -4.0, -0.0, lnNu_g (Probability involved in the generating matrix to go to a selection state from the neutral state at the group level) to -5.0,-0.0, s_max: Maximum state of the Markov model to 4, beta (coefficient of drift of a population) to a mean of -2.0 and standard deviation of 1.8, alpha_max (maximum coefficient of selection of a group) = 4.0, lnKappa (logarithm of group positive scaling parameter) of a minimum of -10.0 and maximum of -0.1, sigmaProp_mu (Value to determine the range of the proposal value of mu) to 0.005, sigmaProp_nu (Value to determine the range of the proposal value of mu) to 0.05, sigmaProp_kappa (Value to determine the range of the proposal value of lnkappa) to 0.05. The number of iterations was set to 500,000, the burn-in to 300,000 and the thinning to 100.

To get further insights into the results obtained by Flink, we plotted the raw genotypes of all the SNPs, SNPs under homogenising selection in the coastal populations, SNPs under divergent selection between ecotypes, and SNPs under both homogenising selection in the coastal populations and divergent between ecotypes (defined as the SNPs under parallel selection) using the R packages vcfR (52) and adegenet (53, 54). We also plotted a neighbor-joining distance tree for the SNPs under each type of selection using the R package ape (55) and a PCA using the packages adegenet (glPCA function) and scales (42). To determine the origin of the SNPs under selection, we defined how many were also polymorphic in the pelagic populations, and compared the 2DSFS between all pairs of populations, estimated in ANGSD (see details in Supplementary text), using all SNPs, the SNPs under homogenising selection in the coastal populations and SNPs under divergent selection between ecotypes. Then, we defined how many SNPs under the two types of selection and under parallel selection were found in ancient tracts. We compared the results with 100 random samples of the same number of putatively neutral SNPs found in ancient tracts.

Putative functions of the genes under selection were determined using literature search, Entrez Gene (56), Uniprot (57), RefSeq (58), GeneCards(59) and Online Mendelian Inheritance in Man (OMIM)(60) databases.

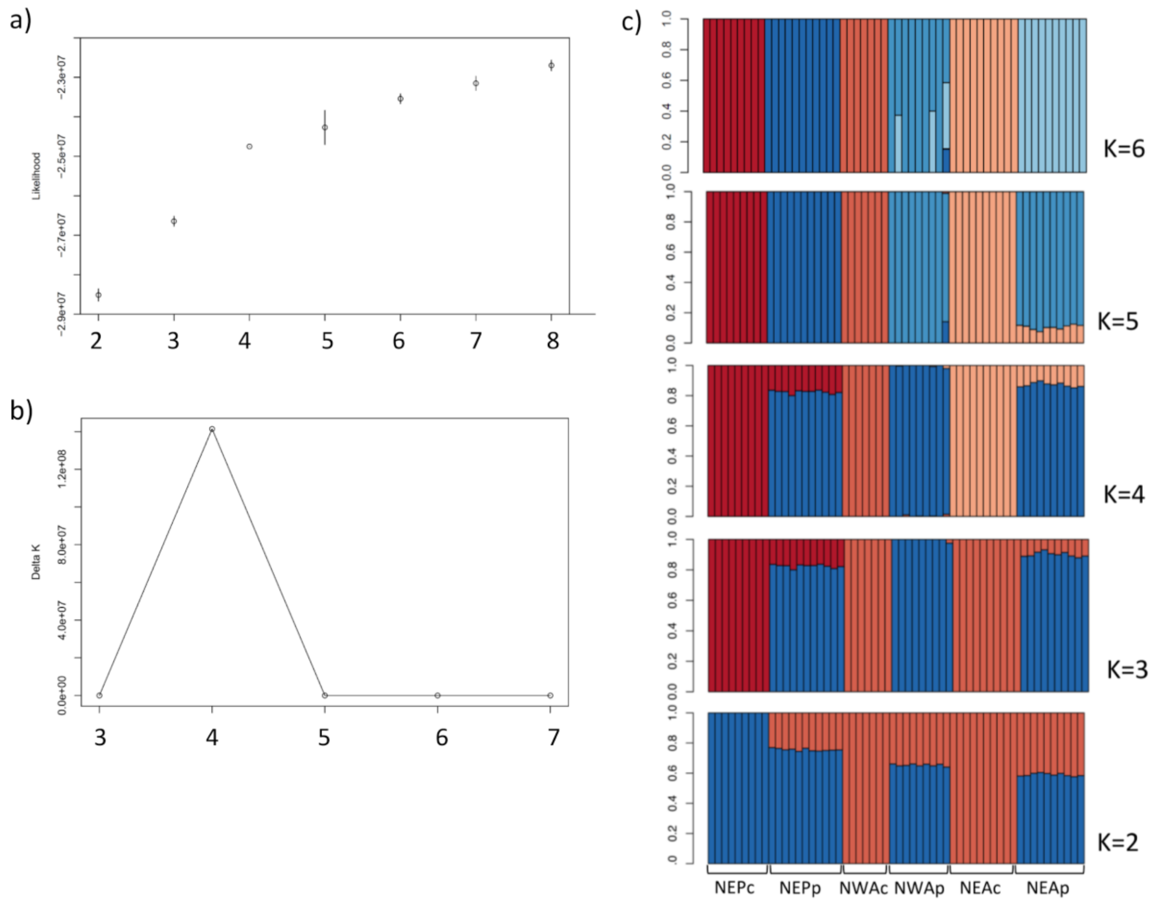


**Supplementary figure 1.** a) NGSAdmix log likelihoods and b) the rate of likelihood change (Delta *K*) (Evanno et al. 2005) for each number of cluster (K) values. c) Ancestry proportions for each of the 57 individuals inferred in NGSAdmix for K=2 to K=6.

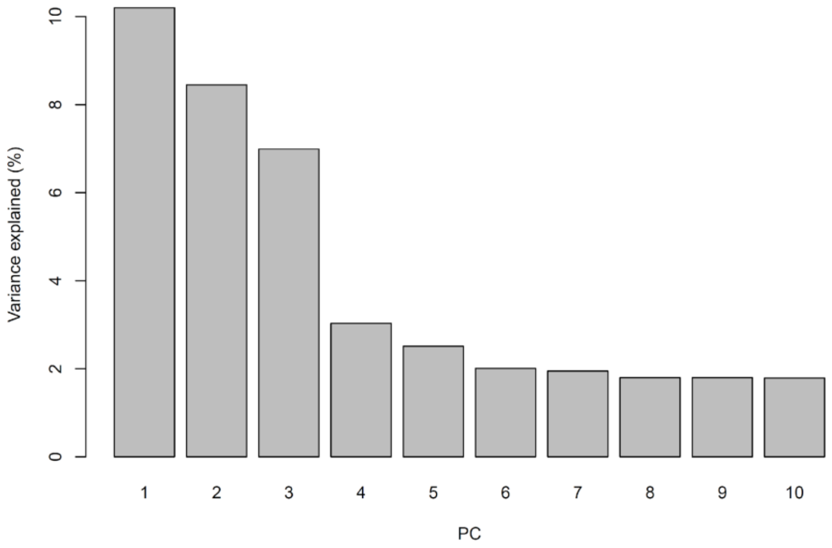


**Supplementary figure 2.** Principal component analysis scree plot showing the percentage of variance explained by each of the 10 first principal components (PCs).

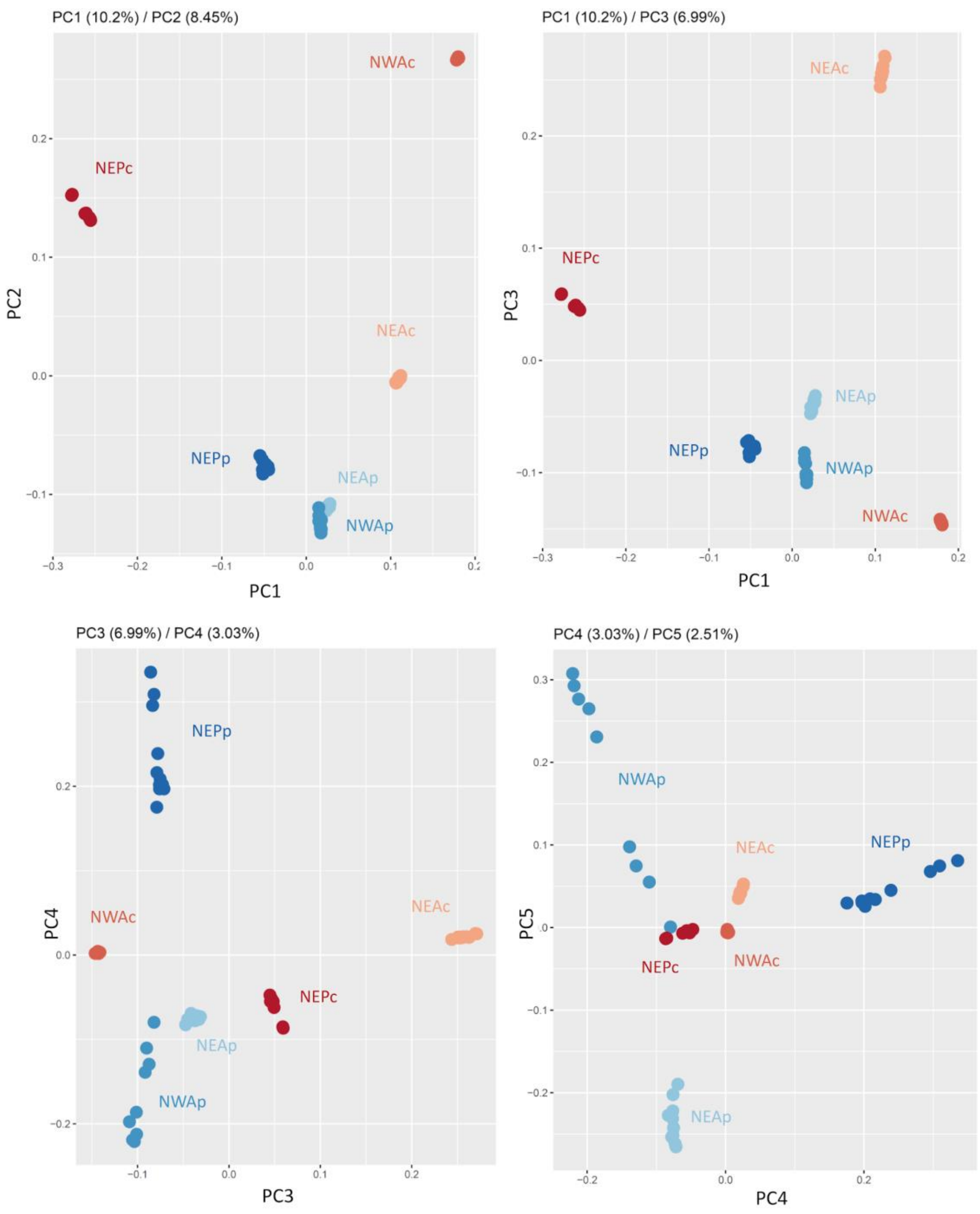


**Supplementary figure 3.** Principal component analysis showing the first and second, first and third, third and fourth, and fourth and fifth principal components (PCs).

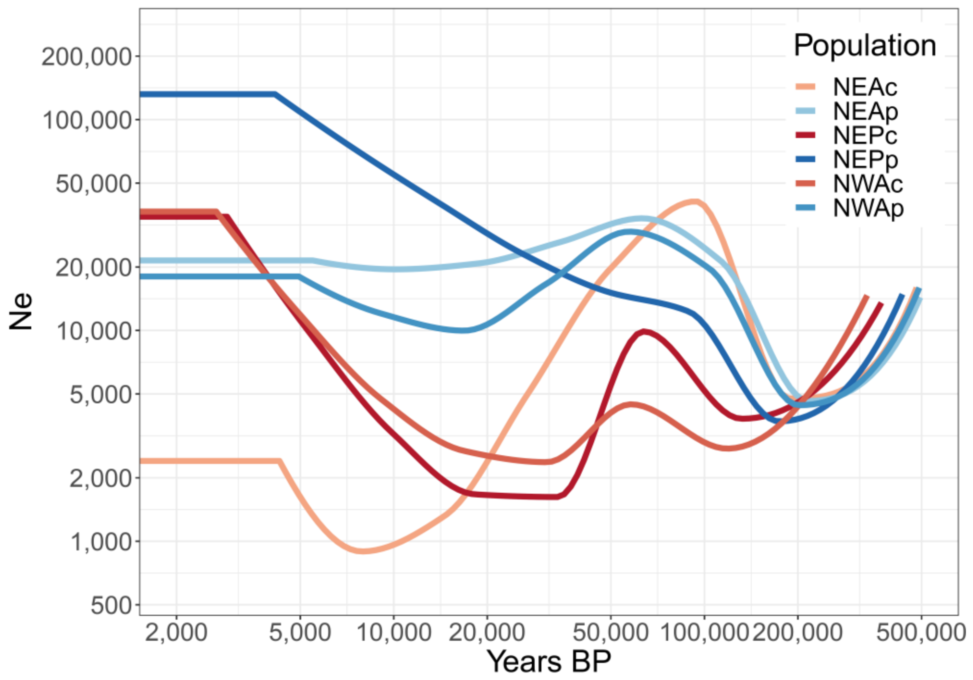


**Supplementary figure 4.** Changes in effective population size through time inferred for each bottlenose dolphin population using SMC++ using a mutation rate of 1.92e-8 substitution per nucleotide per generation (Dornburg et al. 2012) and a generation time of 21.1 years (Taylor et al. 2007). Only neutral sites are included.

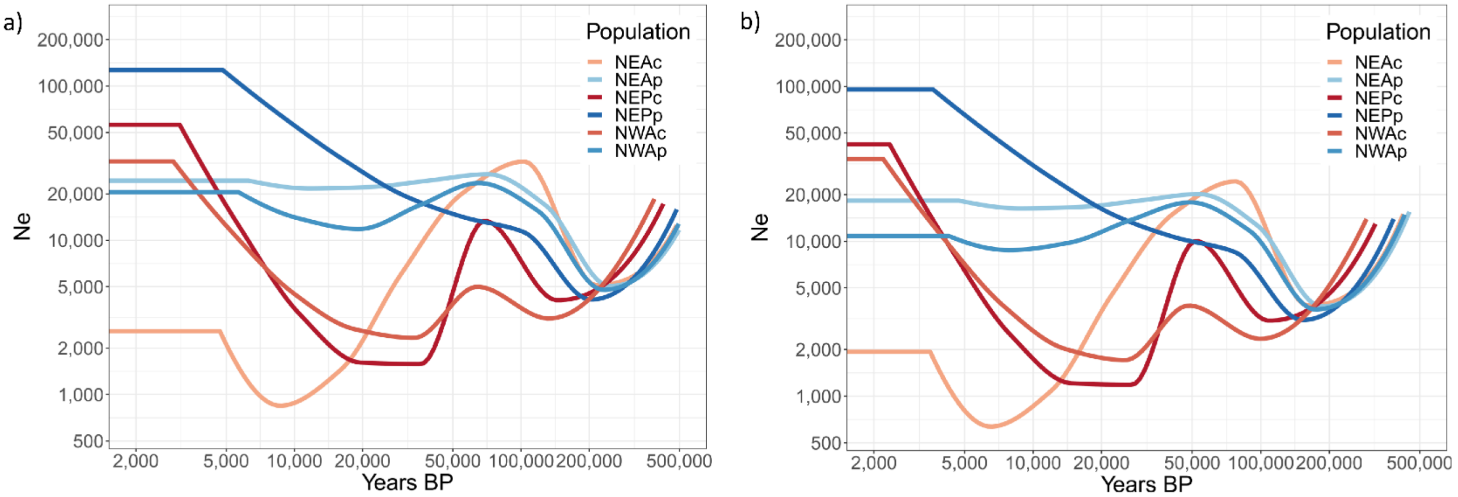


**Supplementary figure 5.** Changes in effective population size through time inferred for each bottlenose dolphin population for all sites (neutral and under selection as identified by Flink) using SMC++, a generation time of 21.1 years (Taylor et al. 2007), and a) a mutation rate of 1.92e-8 substitution per nucleotide per generation (Dornburg et al. 2012) and b) a mutation rate of 2.56e-8 substitution per nucleotide per generation (Yim et al. 2014).

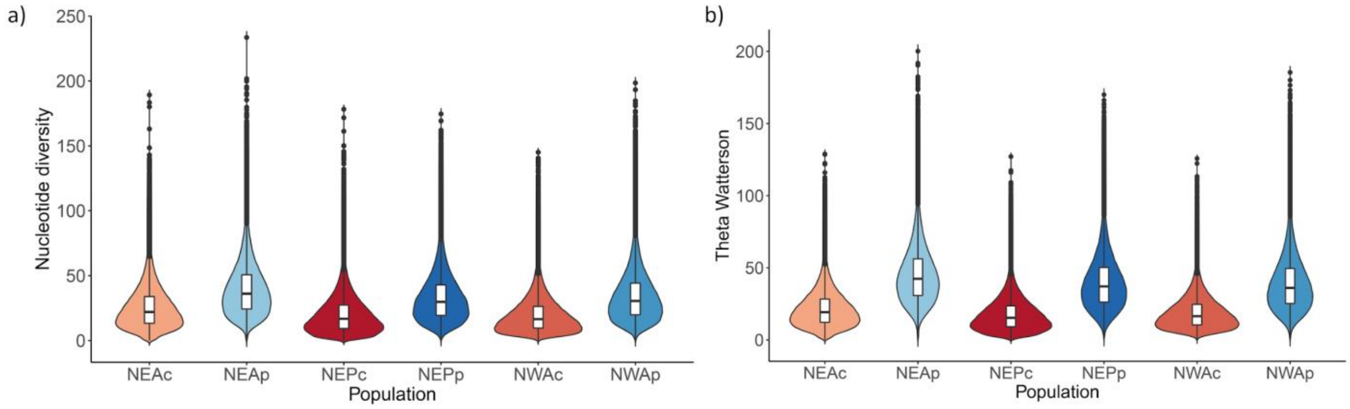


**Supplementary figure 6.** A) Nucleotide diversity and b) Theta Watterson estimated for each population of bottlenose dolphins from the SFS (Supplementary figure 12).

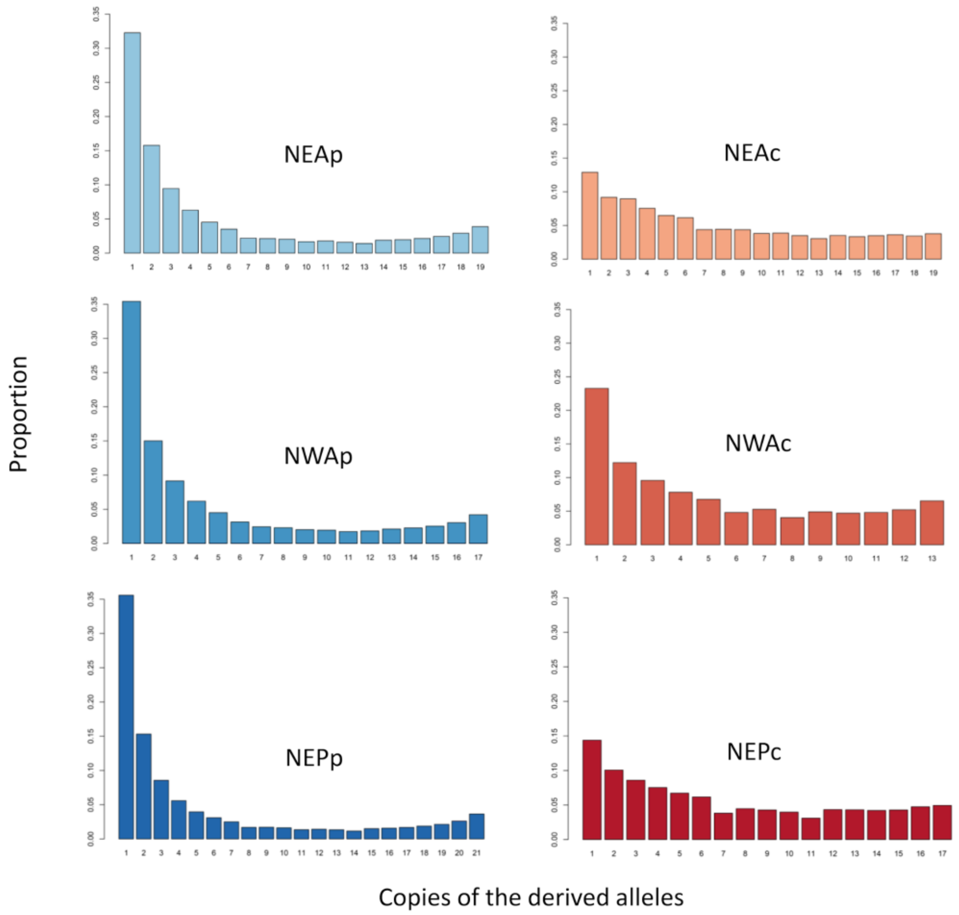


**Supplementary Figure 7.** Site-Frequency-Spectrum for each of the bottlenose dolphin population, as the proportion of each number of copies of the derived alleles.

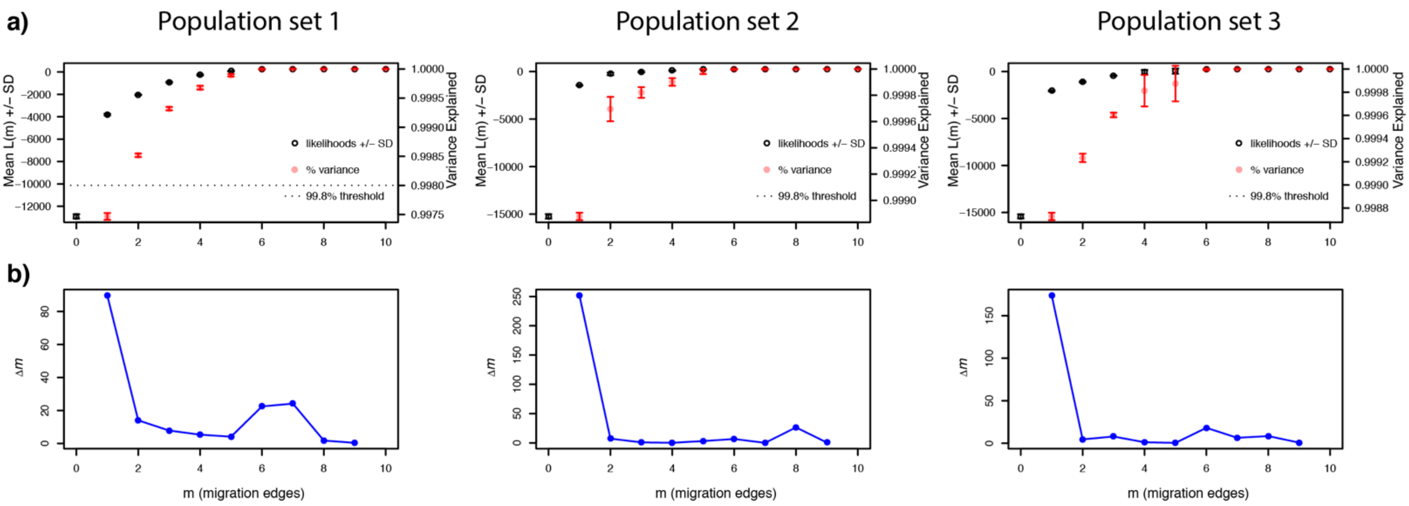


**Supplementary figure 8.** Determination of the optimal number of migration edges between 0 and 10 in TreeMix using the a) log-likelihood values and percentage of variance explained, and b) the second order rate of change in the log-likelihood (Evanno) method for the three sets of individuals.

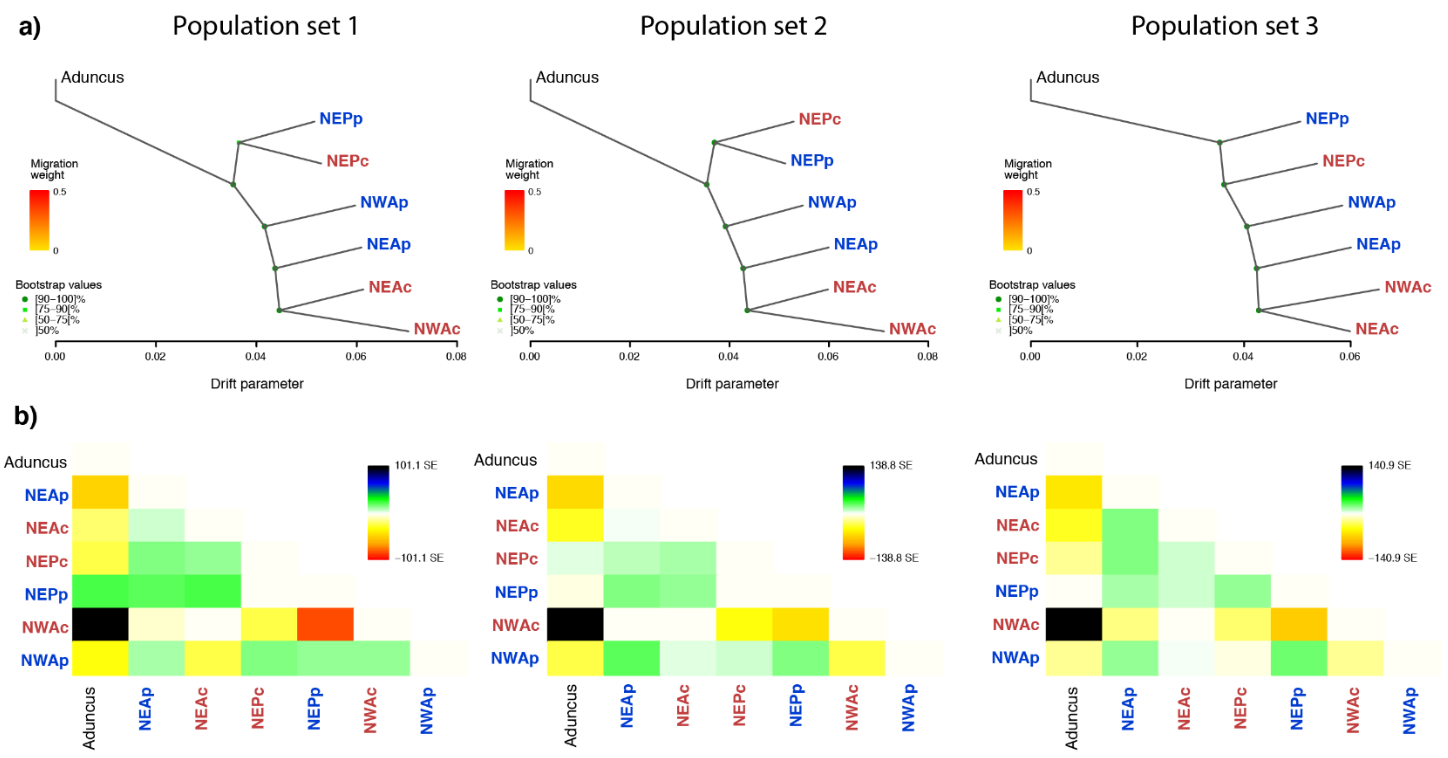


**Supplementary Figure 9**. a) TreeMix consensus tree and bootstrap values displaying the relationships among populations as a bifurcating maximum-likelihood tree for no migration edges for the three sets of individuals. Horizontal branch lengths represent the amount of genetic drift that has occurred along each branch. b) Residual fit of the observed versus the predicted squared allele frequency difference, expressed as the number of SE of the deviation for no migration. SE values are represented by colours according to the palette on the right. Residuals above zero indicate populations that are more closely related to each other in the data than in the best-fit tree and have potentially undergone admixture. Negative residuals represent populations that are less closely related in the data than represented in the best-fit tree.

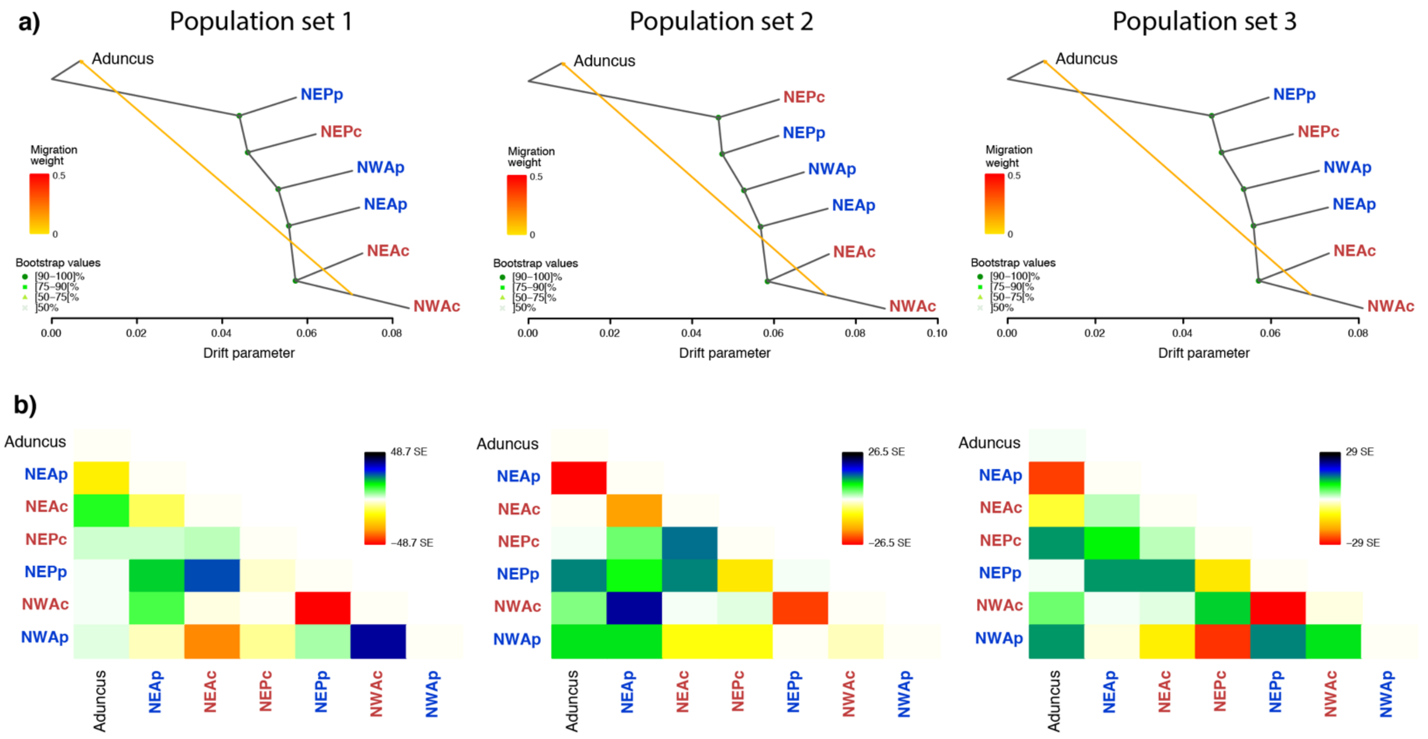


**Supplementary Figure 10.** a) TreeMix consensus tree and bootstrap values displaying the relationships among populations as a bifurcating maximum-likelihood tree with one migration edge for the three sets of individuals. Horizontal branch lengths represent the amount of genetic drift that has occurred along each branch. b): Residual fit of the observed versus the predicted squared allele frequency difference, expressed as the number of SE of the deviation with one migration edge. SE values are represented by colours according to the palette on the right. Residuals above zero indicate populations that are more closely related to each other in the data than in the best-fit tree and have potentially undergone admixture. Negative residuals represent populations that are less closely related in the data than represented in the best-fit tree.

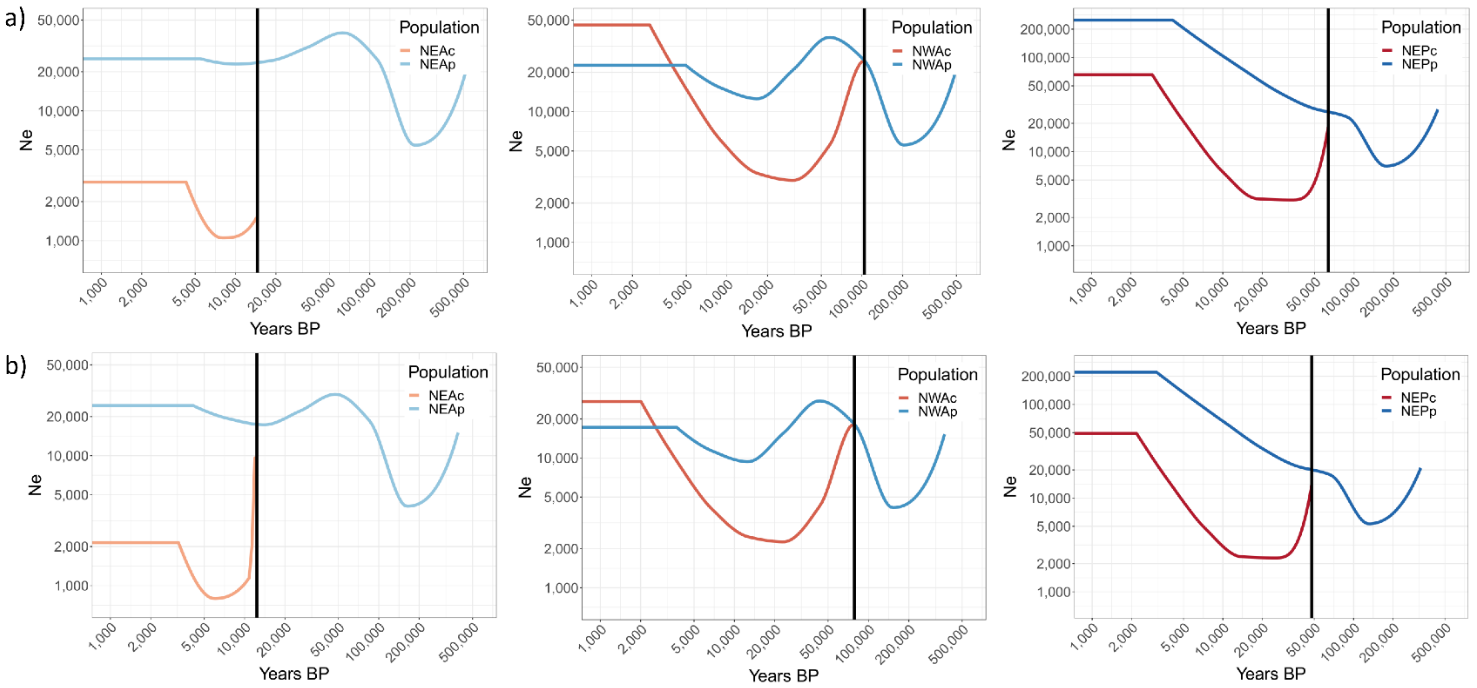


**Supplementary figure 11**. Divergence time between ecotype pairs in the NEA, NWA and NEP estimated using SMC++, a generation time of 21.1 years (Taylor et al. 2007), and a) a mutation rate of 1.92e-8 substitution per nucleotide per generation (Dornburg et al. 2012) and b) a mutation rate of 2.56e-8 substitution per nucleotide per generation (Yim et al. 2014).

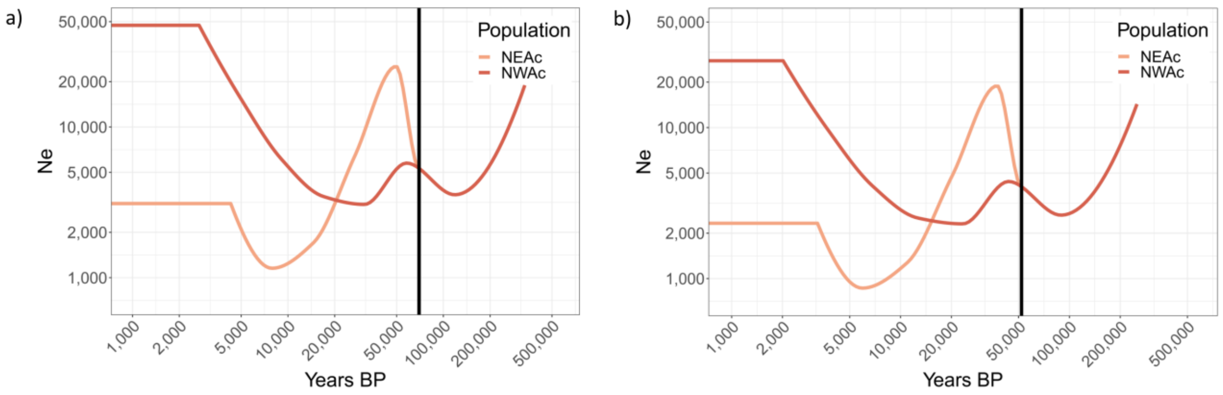


**Supplementary figure 12.** Divergence time between the two coastal populations in the North Atlantic estimated using SMC++, a generation time of 21.1 years (Taylor et al. 2007), and a) a mutation rate of 1.92e-8 substitution per nucleotide per generation (Dornburg et al. 2012) and b) a mutation rate of 2.56e-8 substitution per nucleotide per generation (Yim et al. 2014).

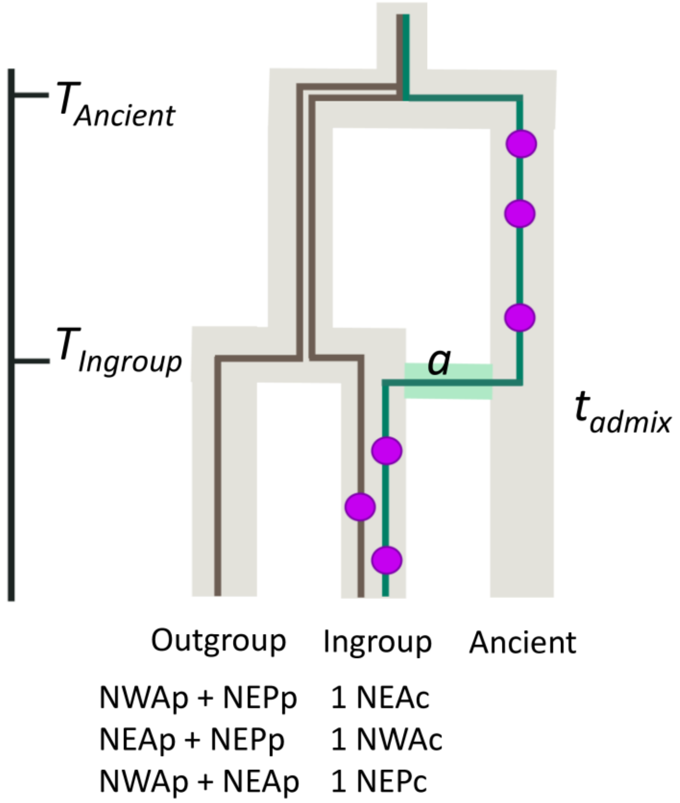


**Supplementary figure 13.** Ancient ancestry method's principles, re-drawn from Skov et al. 2018. At time $T_{admix}$, an ancient tract introgresses from a ghost population into the ingroup population, here one of the coastal population, with admixture proportion $a$. We test for ancient introgressions into each coastal population. All non-allopatric pelagic samples are part of the outgroup. The method scans the genome for cluster of private alleles (purple circles) in each coastal population. Ancient introgressed regions will have higher private variants density than non-introgressed. This is because the split between the pelagic populations and the ghost population – $T_{Ancient}$ – is older than the split between the pelagic and coastal populations $T_{Ingroup}$. Therefore, ancient tracts have had more time to accumulate variation not found in the pelagic populations. However, we hypothesis that ancient tracts in coastal bottlenose dolphin populations may not have been directly introgressed but rather have been retained as balanced polymorphism, see discussion.

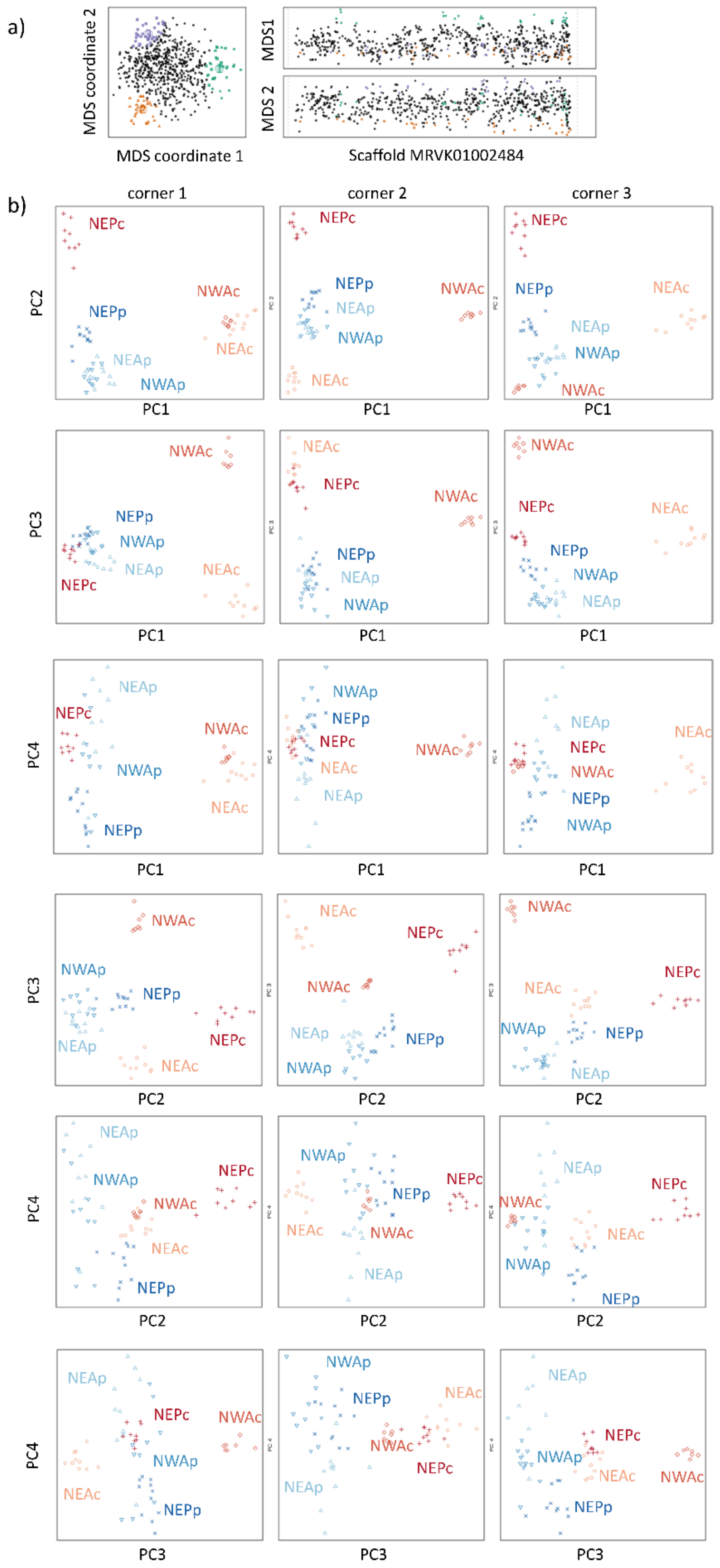


**Supplementary figure 14.** Local PCA results. a) MDS identifying the three major patterns of relatedness among bottlenose dolphin populations and b) PCA describing the three major patterns of relatedness (corners 1-3, green, orange and purple respectively) among populations on four PCs for scaffold MRVK01002484.

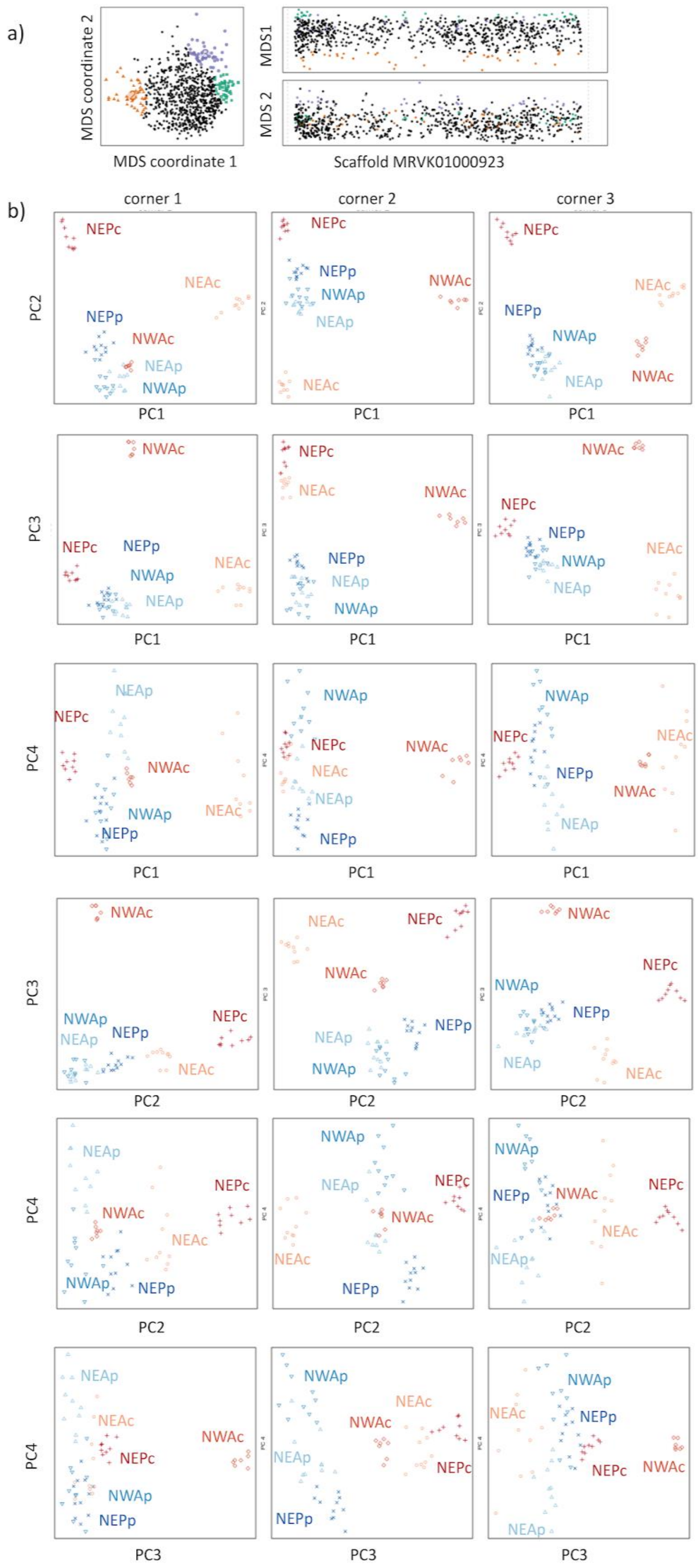


**Supplementary figure 15.** Local PCA results. a) MDS identifying the three major patterns of relatedness among bottlenose dolphin populations and b) PCA describing the three major patterns of relatedness (corners 1-3, green, orange and purple respectively) among populations on four PCs for scaffold MRVK01000923.

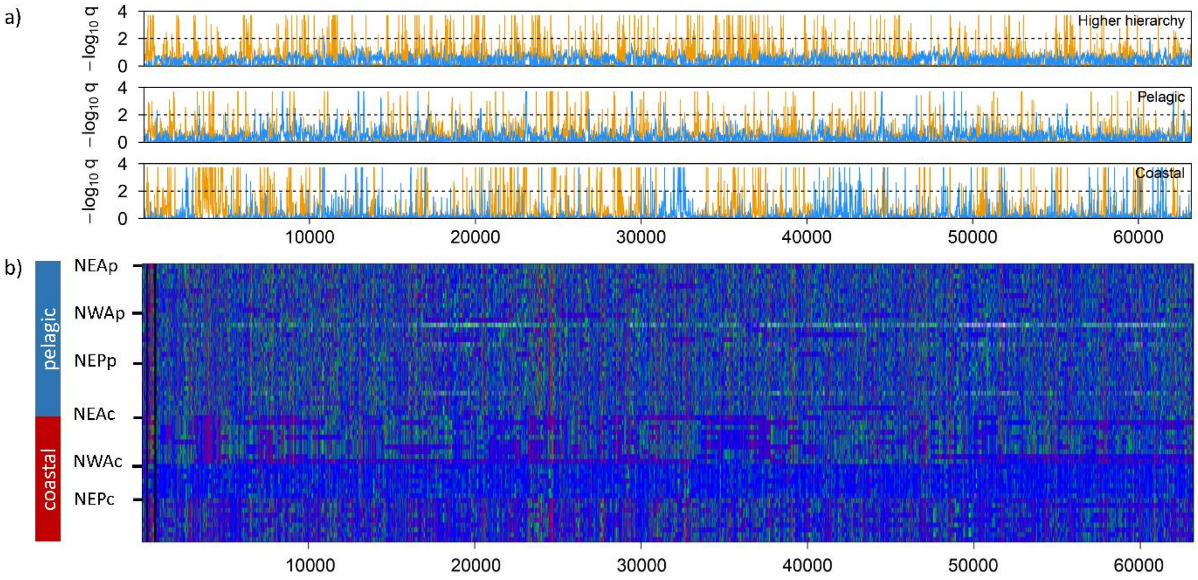


**Supplementary figure 16.** a) Patterns of selection (divergent: yellow, homogenising: blue) inferred using Flink from one scaffold grouping between coastal and pelagic population (top panel), among pelagic populations (middle panel) and among coastal populations (lower panel) for scaffold ensemble 4. The y-axis indicate the locus-specific FDR for divergent (orange) and homogenising (blue) selection, respectively. The black dashed line shows the 1% FDR threshold, above which we consider the locus under selection. b) Plot of the genotypes along scaffold 4, with blue: homozygote reference, green: heterozygote, and red: homozygote alternated. Black lines indicate the separation between different scaffolds.

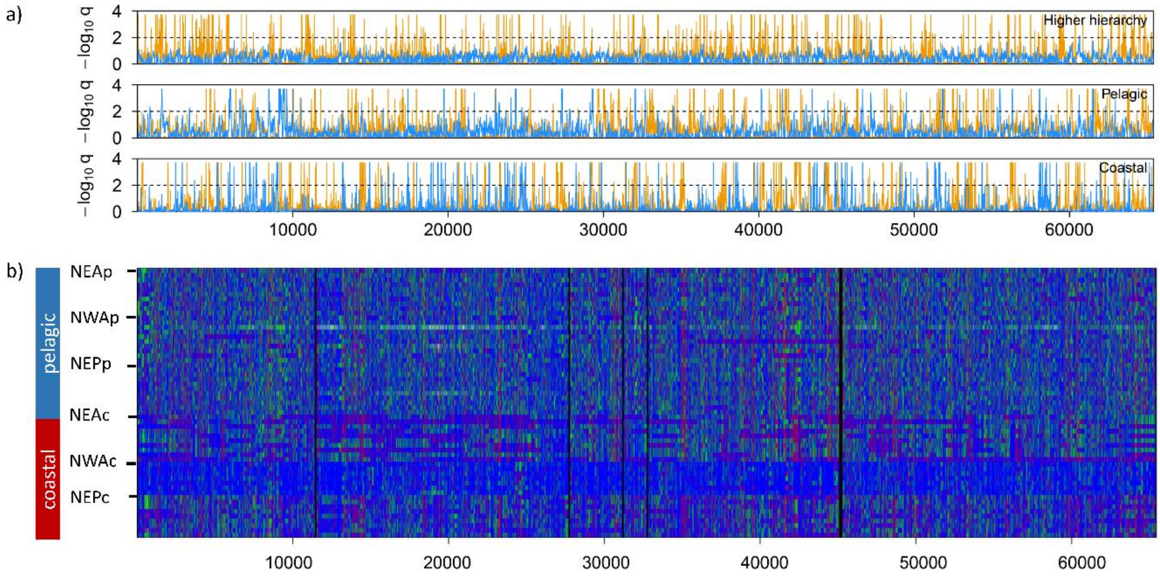


**Supplementary figure 17.** a) Patterns of selection (divergent: yellow, homogenising: blue) inferred using Flink from one scaffold grouping between coastal and pelagic population (top panel), among pelagic populations (middle panel) and among coastal populations (lower panel) for scaffold ensemble 9. The y-axis indicate the locus-specific FDR for divergent (orange) and homogenising (blue) selection, respectively. The black dashed line shows the 1% FDR threshold, above which we consider the locus under selection. b) Plot of the genotypes along scaffold 9, with blue: homozygote reference, green:

heterozygote, and red: homozygote alternated. Black lines indicate the separation between different scaffolds.

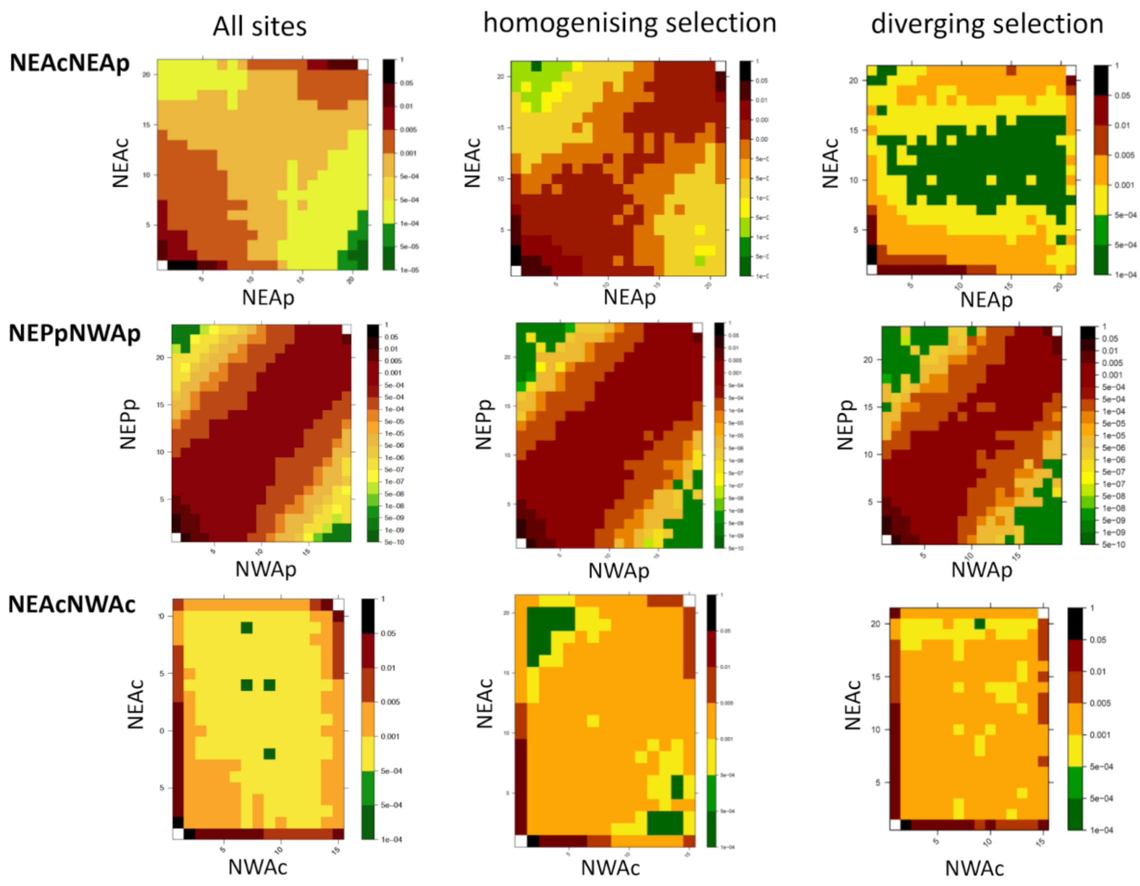


**Supplementary figure 18.** 2D-Site-Frequency-Spectrum (SFS) for all sites (left panels), sites under homogenising selection in the coastal population (middle panels), and sites under divergent selection between ecotypes (right panels) for a coastal and a pelagic population (NEAc-NEAp), two pelagic populations (NEPp-NWAp) and for two coastal populations (NEAc-NWAc). Note that the patterns seen in these populations is similar across all pairs.

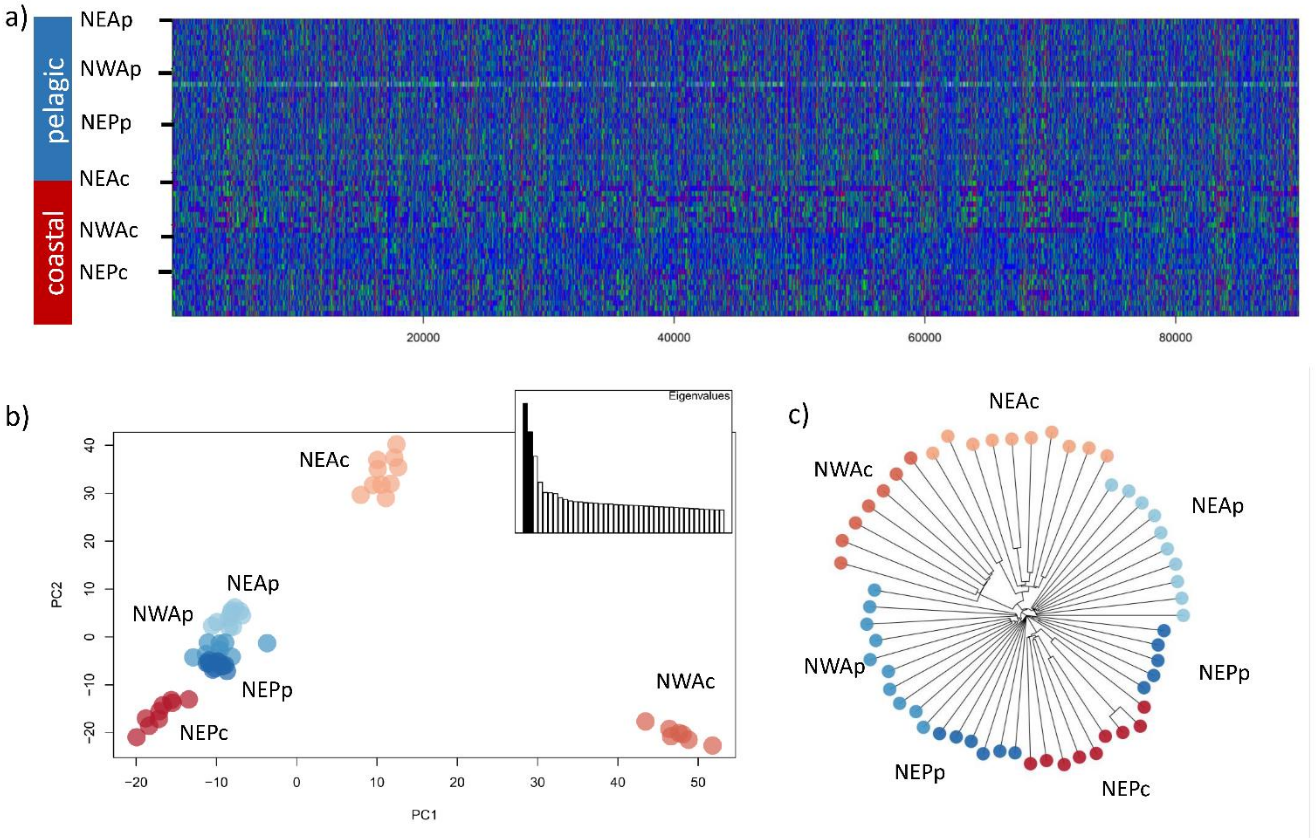


**Supplementary figure 19.** Patterns of genetic variation of the 89,796 SNPs, scattered across the genome, under homogenising selection among the coastal populations. a) Plot of the genotypes, with blue: homozygote reference, green: heterozygote, and red: homozygote derived, b) Principal Component Analysis and b) Neighbor-Joining distance tree of the common bottlenose dolphin samples.

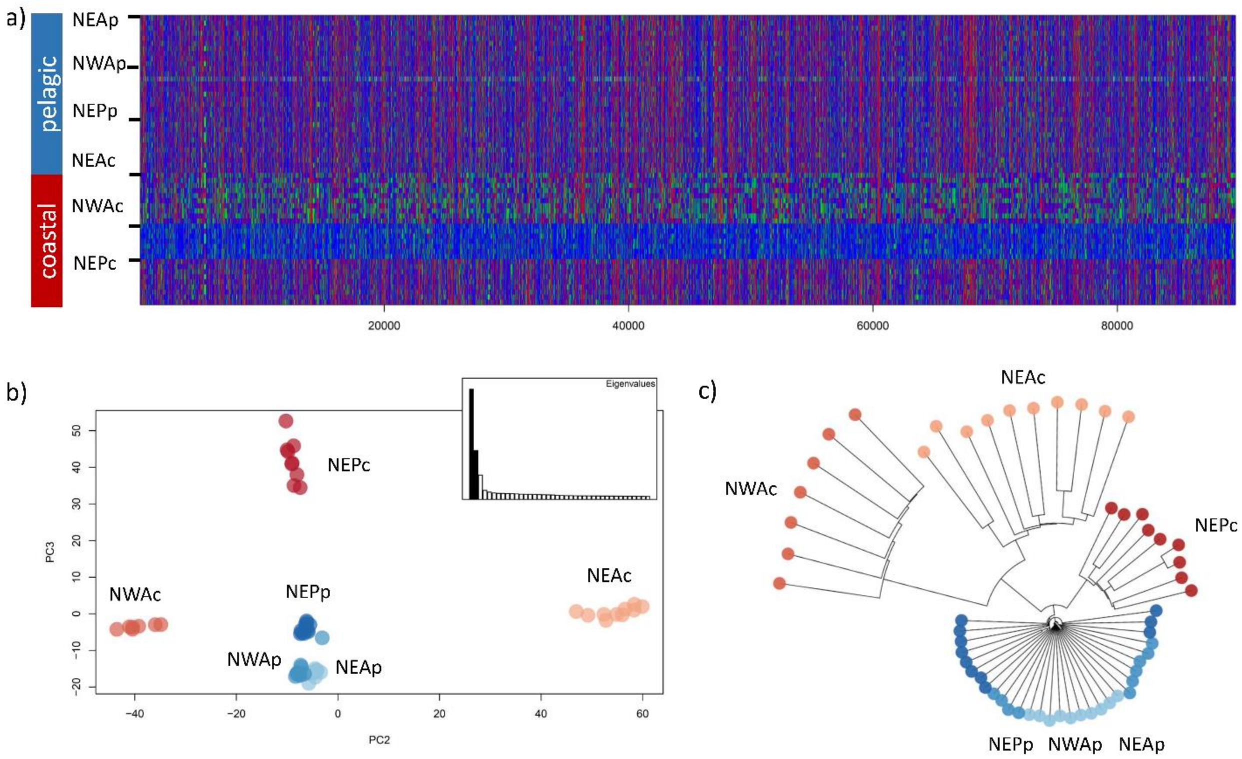


**Supplementary figure 20.** Patterns of genetic variation of the 89,663 SNPs, scattered across the genome, under divergent selection between ecotypes. a) Plot of the genotypes, with blue: homozygote reference, green: heterozygote, and red: homozygote derived, b) Principal Component Analysis and b) Neighbor-Joining distance tree of the common bottlenose dolphin samples.

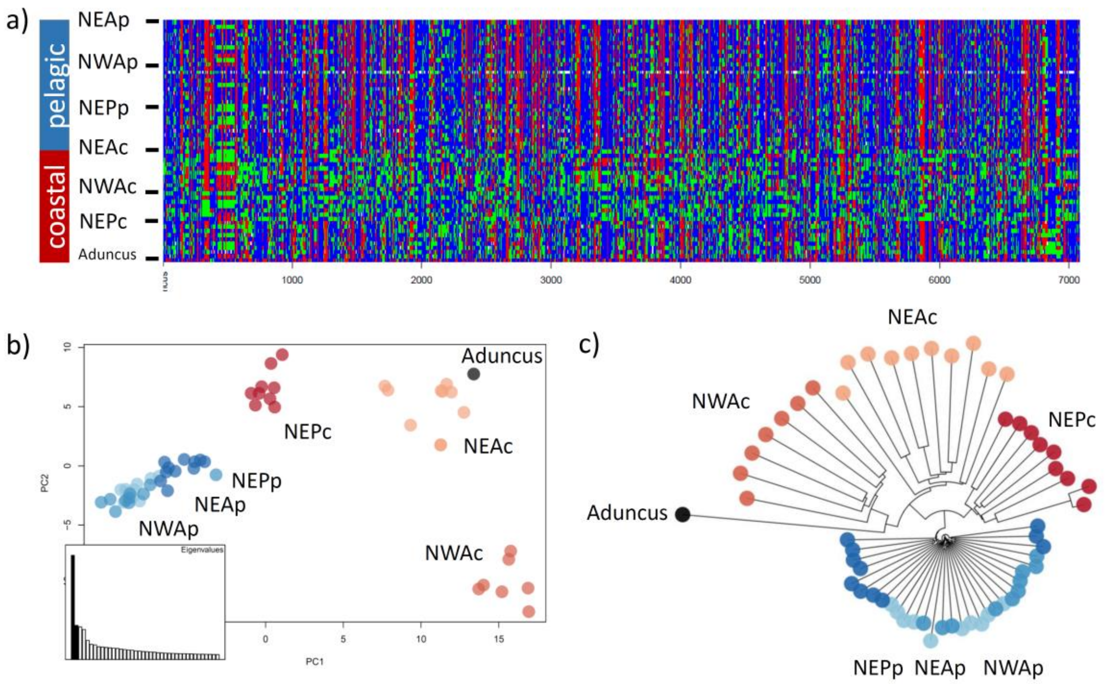


**Supplementary figure 21.** Patterns of genetic variation of the 7,165 SNPs under parallel selection to coastal habitat, i.e. under both homogenising selection among coastal population and divergent selection between ecotypes, including the Indo-Pacific bottlenose dolphin. These SNPs are scattered across the genome. a) Plot of the homozygote reference genotypes in blue, heterozygote in green and homozygote for the alternated allele in red. b) Principal component analysis and (c) Neighbor-joining distance tree showing the genetic structure of the common bottlenose dolphin samples and the Indo-Pacific bottlenose dolphin sample for this particular SNP set.

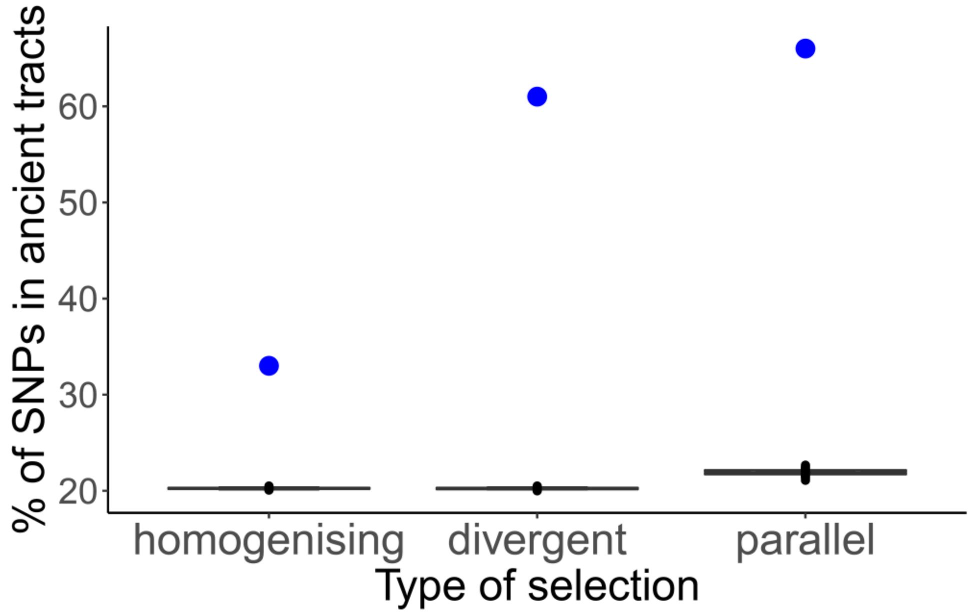


**Supplementary figure 21.** Percentage of SNPs in ancient tracts under the different types of selection: i) homogenising selection among coastal populations “homogenising”, ii) divergent selection between ecotypes “divergent” and iii) in “parallel” selection, that is, both under homogenizing selection among coastal populations and divergent selection between ecotypes, with the blue dots representing the observed percentage in our data and the boxplots the percentage in 100 random sample sets of the same number of putatively neutral SNPs.

**Supplementary table 1.** Sample information indicating the sample ID used in the laboratory (Lab_ID), sample ID used at the institution where the sample was taken (Institute_ID), population, sex, sampling date, number of raw reads after sequencing, mean coverage after all quality filtering including repeats, excessive coverage, mapping quality and phred score, and accession number.

| Lab_ID | Institute_ID | Population | Sex | Sampling date | Number of raw reads | mean coverage after filtering | Accession number |
|---|---|---|---|---|---|---|---|
| S27 | SW1992/201c | NEAc | M | 10/11/1992 | 272014714 | 8.73 | |
| S31 | SW2011/173 | NEAc | M | 4/23/2011 | 254077360 | 8.75 | |
| S37 | SW1995/145a | NEAc | M | 12/31/1995 | 237369566 | 8.44 | |
| S38 | SW1995/99a | NEAc | F | 8/6/1995 | 260063378 | 8.52 | |
| S39 | SW1993/11c | NEAc | M | 1/25/1993 | 231416096 | 7.78 | |
| S41 | SW1996/103c | NEAc | M | 6/17/1996 | 250014606 | 8.66 | |
| S42 | SW2004/257a | NEAc | M | 8/20/2004 | 232352130 | 7.53 | |
| S53 | SW1999/66b | NEAc | F | 3/30/1999 | 256252676 | 8.78 | |
| S54 | SW1999/136A | NEAc | F | 7/12/1999 | 265370600 | 8.80 | |
| S56 | SW2001/111a | NEAc | F | 5/23/2001 | 244673284 | 8.23 | |
| IR23 | 2007.1.179 | NEAp | M | 12/07/2009 | 282095448 | 9.13 | |
| IR24 | 2007.1.180 | NEAp | M | 21/07/2009 | 278318284 | 9.34 | |
| IR30 | 2007.1.276 | NEAp | M | 10/04/2012 | 236082662 | 8.63 | |
| IR33 | 2007.1.273 | NEAp | F | 17/05/2011 | 309762602 | 10.46 | |
| IR34 | 2007.1.272 | NEAp | F | 27/01/2012 | 231376318 | 8.07 | |
| IR36 | 2007.1.270 | NEAp | M | 03/06/2011 | 260728180 | 8.80 | |
| S10 | SW2001/75a | NEAp | M | 4/2/2001 | 337738358 | 11.32 | |
| S12 | SW1998/18a | NEAp | F | 1/25/1998 | 235381406 | 7.75 | |
| S40 | SW2007/4c | NEAp | M | 1/6/2007 | 298305240 | 9.95 | |
| S43 | SW2011/188 | NEAp | M | 5/1/2011 | 227419788 | 7.94 | |
| 115669 | Tt1 | NEPc | M | 26/6/2013 | 270809186 | 6.53 | |
| 117692 | CTTSD131023.01 | NEPc | F | 23/10/2013 | 267379282 | 6.17 | |
| 125942 | CTTSD100723.02 | NEPc | M | 23/7/2010 | 286417592 | 6.89 | |
| 146413 | CTTSD131213.01 | NEPc | M | 13/12/2013 | 264961758 | 6.46 | |
| 146414 | CTTSD131213.02 | NEPc | M | 13/12/2013 | 282157544 | 6.83 | |
| 146416 | HYDE150113.01 | NEPc | M | 13/1/2015 | 272176732 | 6.53 | |
| 146419 | HYDE150128.01 | NEPc | F | 28/1/2015 | 271565530 | 6.46 | |
| 160317 | LSK151102.04 | NEPc | M | 2/11/2015 | 216981988 | 5.33 | |
| 92200 | CTTSD091102.01 | NEPc | M | 2/11/2009 | 282378954 | 6.52 | |
| 113122 | CTTSD121029.06 | NEPp | M | 29/10/2012 | 266083128 | 6.21 | |
| 117696 | GCAMPBELL131102.02 | NEPp | F | 2/11/2013 | 245179228 | 6.01 | |
| 117698 | GCAMPBELL131103.01 | NEPp | M | 3/11/2013 | 262410774 | 6.59 | |
| 125944 | CTTSD100730.02 | NEPp | M | 30/7/2010 | 274668084 | 6.55 | |
| 145428 | DSJ141023.15 | NEPp | F | 23/10/2014 | 289591618 | 6.59 | |
| 145429 | DSJ141023.16 | NEPp | M | 23/10/2014 | 247400236 | 5.90 | |
| 145430 | DSJ141023.17 | NEPp | F? | 23/10/2014 | 224105208 | 4.30 | |
| 160321 | LSK151104.02 | NEPp | F | 4/11/2015 | 254454974 | 6.04 | |
| 160322 | LSK151104.03 | NEPp | M | 4/11/2015 | 265724004 | 6.38 | |
| 160326 | LSK151105.04 | NEPp | F | 5/11/2015 | 275175338 | 6.41 | |

| | | | | | | |
|---|---|---|---|---|---|---|
| 160335 | LSK151107.06 | NEPp | F | 7/11/2015 | 271266882 | 6.62 |
| 14Tt001 | 173059 | NWAc | M | 14/3/2002 | 310465756 | 9.82 |
| 14Tt004 | 173060 | NWAc | F | 14/3/2002 | 271870888 | 7.51 |
| 14Tt006 | 173061 | NWAc | M | 14/3/2002 | 309447666 | 9.46 |
| 14Tt021 | 173062 | NWAc | M | 16/3/2002 | 292191584 | 9.33 |
| 14Tt022 | 173063 | NWAc | M | 16/3/2002 | 317306584 | 7.74 |
| 14Tt023 | 173064 | NWAc | M | 16/3/2002 | 275133508 | 7.16 |
| 14Tt025 | 173065 | NWAc | F | 16/3/2002 | 304170502 | 6.50 |
| 7Tt156 | 173048 | NWAp | F | 10/8/1999 | 238633104 | 7.27 |
| 7Tt161 | 173049 | NWAp | M | 11/8/1999 | 278592266 | 8.67 |
| 7Tt182* | 173051 | NWAp | M | 16/8/1999 | 222939912 | 2.62* |
| 7Tt193 | 173052 | NWAp | M | 14/8/1999 | 263546918 | 7.73 |
| 7Tt270 | 173053 | NWAp | M | 17/3/2002 | 217855012 | 7.08 |
| 7Tt278 | 173054 | NWAp | M | 1/4/2002 | 260697324 | 8.05 |
| 7Tt282 | 173055 | NWAp | M | 2/4/2002 | 254492318 | 6.02 |
| 7Tt284 | 173056 | NWAp | F | 2/4/2002 | 252067218 | 6.87 |
| 7Tt287 | 173057 | NWAp | F | 3/4/2002 | 244155564 | 8.02 |
| 7Tt350 | 173058 | NWAp | M | 13/7/2005 | 253335024 | 7.91 |

* Due to its low coverage, this individual was excluded from analyses not based on allele frequencies

**Supplementary table 2**. Mean pairwise weighted $F_{ST}$ as estimated in vcftools[92] using a sliding-window size of 50 kb and a step size of 10 kb.

| $F_{ST}$ | NEPc | NEPp | NWAc | NWAp | NEAc | NEAp |
|---|---|---|---|---|---|---|
| NEPc | | 0.15 (0.09) | 0.45 (0.18) | 0.20 (0.10) | 0.34 (0.15) | 0.19 (0.10) |
| NEPp | | | 0.29 (0.15) | 0.04 (0.04) | 0.20 (0.12) | 0.04 (0.04) |
| NWAc | | | | 0.28 (0.15) | 0.36 (0.19) | 0.27 (0.14) |
| NWAp | | | | | 0.20 (0.12) | 0.02 (0.04) |
| NEAc | | | | | | 0.16 (0.11) |
| NEAp | | | | | | |

**Supplementary table 3.** Number of regions identified as ancient tracts with posterior probabilities of >0.8 and their total length as estimated following Skov et al. 2018 (see Supplementary figure 13).

| Individual | number of regions with P>0.8 | length in bp |
|---|---|---|
| NEAc1 | 1,464 | 15,487,000 |
| NEAc2 | 1,172 | 14,910,000 |
| NEAc3 | 1,136 | 14,380,000 |
| NEAc4 | 1,174 | 15,209,000 |
| NEAc5 | 1,243 | 13,937,000 |
| NEAc6 | 1,210 | 12,362,000 |
| NEAc7 | 1,207 | 15,273,000 |
| NEAc8 | 1,339 | 14,320,000 |
| NEAc9 | 1,241 | 14,778,000 |
| NEAc10 | 1,141 | 14,725,000 |
| NWAc1 | 2,067 | 24,861,000 |
| NWAc2 | 1,755 | 22,939,000 |
| NWAc3 | 1,982 | 20,528,000 |
| NWAc4 | 2,097 | 23,085,000 |
| NWAc5 | 2,037 | 23,645,000 |
| NWAc6 | 1,853 | 21,869,000 |
| NWAc7 | 2,108 | 15,377,000 |
| NEPc1 | 1,370 | 11,608,000 |
| NEPc2 | 1,373 | 11,867,000 |
| NEPc3 | 1,537 | 12,400,000 |
| NEPc4 | 1,500 | 10,666,000 |
| NEPc5 | 1,496 | 11,161,000 |
| NEPc6 | 1,330 | 11,429,000 |
| NEPc7 | 1,372 | 11,858,000 |
| NEPc8 | 1,678 | 10,185,000 |
| NEPc9 | 1,496 | 12,714,000 |

**Supplementary table 4.** TMRCA between the coastal individual and the allopatric pelagic individual ($T_{Ingroup}$) and between the introgressed tracts within coastal dolphins and the corresponding genomic regions in the outgroup ($T_{Ancient}$, Supplementary Figure 7) using mutation rates 1: 1.92e-8 substitution per nucleotide per generation (Dornburg et al. 2012), and 2: 2.56e-8 substitution per nucleotide per generation (Yim et al. 2014) in generations and in years using a generation time of 21.1 years (Taylor et al. 2007).

| | GENERATIONS | | YEARS | |
|---|---|---|---|---|
| | ***T*ingroup** | ***T*ancient** | ***T*ingroup** | ***T*ancient** |
| **mutation 1** | | | | |
| NEAc1 | 9,688 | 68,521 | 204,425 | 1,445,793 |
| NEAc2 | 9,384 | 63,602 | 198,007 | 1,342,000 |
| NEAc3 | 9,559 | 61,874 | 201,687 | 1,305,552 |
| NEAc4 | 9,032 | 62,931 | 190,578 | 1,327,853 |
| NEAc5 | 9,447 | 70,583 | 199,336 | 1,489,297 |
| NEAc6 | 9,296 | 71,259 | 196,155 | 1,503,556 |
| NEAc7 | 9,066 | 62,087 | 191,284 | 1,310,043 |
| NEAc8 | 9,373 | 68,912 | 197,760 | 1,454,047 |
| NEAc9 | 9,123 | 64,318 | 192,505 | 1,357,103 |
| NEAc10 | 11,302 | 67,735 | 238,465 | 1,429,215 |
| NWAc1 | 15,564 | 104,506 | 328,405 | 2,205,080 |
| NWAc2 | 18,583 | 107,003 | 392,108 | 2,257,754 |
| NWAc3 | 16,676 | 102,630 | 351,859 | 2,165,498 |
| NWAc4 | 15,004 | 106,478 | 316,580 | 2,246,687 |
| NWAc5 | 15,364 | 100,307 | 324,190 | 2,116,469 |
| NWAc6 | 17,498 | 105,878 | 369,203 | 2,234,025 |
| NWAc7 | 20,115 | 107,629 | 424,417 | 2,270,966 |
| NEPc1 | 10,311 | 76,607 | 217,570 | 1,616,405 |
| NEPc2 | 10,366 | 76,303 | 218,723 | 1,610,002 |
| NEPc3 | 10,610 | 80,668 | 223,869 | 1,702,089 |
| NEPc4 | 10,532 | 81,946 | 222,222 | 1,729,066 |
| NEPc5 | 10,352 | 79,208 | 218,437 | 1,671,298 |
| NEPc6 | 10,312 | 75,117 | 217,582 | 1,584,968 |
| NEPc7 | 10,454 | 78,553 | 220,582 | 1,657,478 |
| NEPc8 | 10,677 | 80,468 | 225,279 | 1,697,869 |
| NEPc9 | 10,380 | 76,218 | 219,028 | 1,608,199 |
| **mutation 2** | | | | |
| NEAc1 | 4,592 | 32,474 | 96,884 | 685,210 |
| NEAc2 | 4,448 | 30,143 | 93,842 | 636,019 |
| NEAc3 | 4,530 | 29,324 | 95,586 | 618,745 |
| NEAc4 | 4,281 | 29,825 | 90,322 | 629,314 |
| NEAc5 | 4,477 | 33,452 | 94,472 | 705,828 |
| NEAc6 | 4,406 | 33,772 | 92,964 | 712,586 |

| | | | | |
|---|---|---|---|---|
| NEAc7 | 4,296 | 29,425 | 90,656 | 620,874 |
| NEAc8 | 4,442 | 32,660 | 93,725 | 689,122 |
| NEAc9 | 4,324 | 30,482 | 91,234 | 643,177 |
| NEAc10 | 5,356 | 32,102 | 113,017 | 677,353 |
| NWAc1 | 7,376 | 49,529 | 155,642 | 1,045,062 |
| NWAc2 | 8,807 | 50,712 | 185,833 | 1,070,026 |
| NWAc3 | 7,903 | 48,640 | 166,758 | 1,026,302 |
| NWAc4 | 7,111 | 50,464 | 150,038 | 1,064,781 |
| NWAc5 | 7,282 | 47,539 | 153,645 | 1,003,066 |
| NWAc6 | 8,293 | 50,179 | 174,978 | 1,058,780 |
| NWAc7 | 9,533 | 51,009 | 201,146 | 1,076,287 |
| NEPc1 | 4,887 | 36,307 | 103,114 | 766,069 |
| NEPc2 | 4,913 | 36,163 | 103,660 | 763,034 |
| NEPc3 | 5,028 | 38,231 | 106,099 | 806,677 |
| NEPc4 | 4,991 | 38,837 | 105,319 | 819,463 |
| NEPc5 | 4,906 | 37,540 | 103,524 | 792,084 |
| NEPc6 | 4,887 | 35,600 | 103,120 | 751,170 |
| NEPc7 | 4,955 | 37,229 | 104,541 | 785,535 |
| NEPc8 | 5,060 | 38,136 | 106,767 | 804,677 |

**Supplementary table 5.** Percentage of shared ancient tracts between pairs of individuals, as the percentage of ancient tracts in individual 1 (i1) that are ancient as well in individual 2 (i2, above diagonal), and the percentage of ancient tracts in individual 2 that are also ancient in individual 1 (below diagonal). Due to the action of recombination, we do not expect individuals within a population to share 100% of ancient tracts, as windows are small (1 Kb) and we only kept those with a certain posterior probability. In addition, it is not expected, for instance there are few examples of fixed or high frequency ancient hominin tracts within any given modern human population.

| i1\ i2 | NEAc1 | NEAc2 | NEAc3 | NEAc4 | NEAc5 | NEAc6 | NEAc7 | NEAc8 | NEAc9 | NEAc10 | NWAc1 | NWAc2 | NWAc3 | NWAc4 | NWAc5 | NWAc6 | NWAc7 | NEPc1 | NEPc2 | NEPc3 | NEPc4 | NEPc5 | NEPc6 | NEPc7 | NEPc8 | NEPc9 |
|---|---|---|---|---|---|---|---|---|---|---|---|---|---|---|---|---|---|---|---|---|---|---|---|---|---|---|
| **NEAc1** | | 32.8 | 30.4 | 29.1 | 26.1 | 27 | 32.7 | 32.2 | 34.3 | 37.6 | 9.5 | 9.2 | 8.9 | 8.7 | 9.6 | 9.3 | 8.1 | 2.3 | 2.2 | 2.3 | 2.2 | 2.3 | 2.1 | 2.3 | 2.2 | 2.6 |
| **NEAc2** | 34.4 | | 35.9 | 32.7 | 31.5 | 28.2 | 34.1 | 31.4 | 32.7 | 30.7 | 10.1 | 9.5 | 8.9 | 9.4 | 9.9 | 9.2 | 8.2 | 1.9 | 2.2 | 2.1 | 2.2 | 2.1 | 2 | 2 | 1.9 | 2.6 |
| **NEAc3** | 30.1 | 33.8 | | 31.6 | 29.9 | 29.9 | 31.2 | 29.7 | 33.8 | 27.8 | 10.1 | 9.9 | 9.4 | 9.4 | 10 | 9.6 | 8.7 | 2.1 | 2.4 | 2.4 | 2.4 | 2.2 | 2.4 | 2.2 | 2.1 | 2.6 |
| **NEAc4** | 29.6 | 31.7 | 32.5 | | 31.1 | 29.2 | 34.4 | 29.5 | 33.7 | 30.4 | 10.8 | 11.3 | 10.3 | 9.9 | 11.2 | 10.5 | 9 | 2.1 | 2.3 | 2.4 | 2.2 | 2.2 | 2.3 | 2.1 | 2.1 | 2.8 |
| **NEAc5** | 30.5 | 35.1 | 35.3 | 35.8 | | 32.7 | 39.8 | 35.4 | 35.1 | 27.6 | 12 | 11.5 | 10.7 | 10.7 | 12 | 11.3 | 9.6 | 2.2 | 2.3 | 2.4 | 2.6 | 2.3 | 2.4 | 2.3 | 2.1 | 2.6 |
| **NEAc6** | 34 | 35.1 | 39.4 | 37.4 | 36.4 | | 36.7 | 33.7 | 35.4 | 31.5 | 12.1 | 11.7 | 11.3 | 11.4 | 12.4 | 11.5 | 10 | 1.9 | 3 | 2.5 | 2.2 | 2.3 | 2.4 | 2.7 | 2.2 | 2.7 |
| **NEAc7** | 32.4 | 32.2 | 31.3 | 33.5 | 33.7 | 27.8 | | 32.6 | 32.7 | 27.5 | 9.7 | 9.9 | 9.1 | 9.7 | 9.9 | 9.5 | 8.5 | 2.1 | 2.1 | 2.3 | 2.1 | 2.3 | 2 | 2.1 | 2.2 | 2.6 |
| **NEAc8** | 36.7 | 34.1 | 34.3 | 33.1 | 34.5 | 29.5 | 37.5 | | 46.3 | 31.7 | 10.7 | 10.6 | 10.1 | 10.5 | 10.8 | 10.6 | 9.2 | 2.1 | 2.2 | 2.4 | 2.1 | 2.4 | 2.4 | 2.4 | 2.1 | 2.8 |
| **NEAc9** | 35.6 | 32.4 | 33.9 | 34.5 | 31.1 | 28.2 | 34.3 | 42.2 | | 31.4 | 10.4 | 10.1 | 9.6 | 9.6 | 10.3 | 9.9 | 8.8 | 2 | 2.5 | 2.3 | 2.3 | 2.4 | 2 | 2.4 | 2.3 | 2.7 |
| **NEAc10** | 37.7 | 29.4 | 28.2 | 30 | 23.6 | 24.3 | 27.9 | 27.9 | 30.3 | | 9.6 | 9.4 | 9.5 | 9.3 | 9.6 | 9.6 | 8.5 | 2.3 | 2.5 | 2.3 | 2.2 | 2.1 | 2.3 | 2.4 | 2.3 | 2.5 |
| **NWAc1** | 7.6 | 7.7 | 8.2 | 8.5 | 8.2 | 7.4 | 7.9 | 7.6 | 8 | 7.7 | | 51.8 | 49.1 | 52 | 53.7 | 49.9 | 40.8 | 2 | 2.1 | 2.1 | 2 | 2 | 2.1 | 2.2 | 2.1 | 2.3 |
| **NWAc2** | 7.5 | 7.4 | 8.1 | 9.1 | 8 | 7.4 | 8.1 | 7.6 | 7.9 | 7.7 | 52.6 | | 48.3 | 50.7 | 52.4 | 48.6 | 39.8 | 2 | 2.2 | 2.2 | 2 | 2.1 | 2.1 | 2.3 | 2.3 | 2.5 |
| **NWAc3** | 7.2 | 6.9 | 7.8 | 8.3 | 7.5 | 7.1 | 7.5 | 7.2 | 7.5 | 7.7 | 50 | 48.3 | | 49 | 49.8 | 47.1 | 40.4 | 2.1 | 2.1 | 2.2 | 2.2 | 2.1 | 2.1 | 2.4 | 2.2 | 2.4 |
| **NWAc4** | 7.6 | 7.8 | 8.3 | 8.4 | 8 | 7.6 | 8.5 | 8.1 | 8 | 8.1 | 56.3 | 54 | 52.2 | | 56.3 | 51.8 | 42.9 | 2 | 1.9 | 2 | 1.9 | 1.8 | 1.9 | 2 | 2.2 | 2.3 |
| **NWAc5** | 7.5 | 7.3 | 7.9 | 8.5 | 7.9 | 7.4 | 7.7 | 7.4 | 7.7 | 7.4 | 51.7 | 49.7 | 47.2 | 50.1 | | 47.7 | 40 | 1.9 | 2.1 | 2.1 | 1.9 | 2 | 2 | 2.2 | 2.1 | 2.3 |
| **NWAc6** | 8.2 | 7.7 | 8.5 | 9.1 | 8.4 | 7.7 | 8.4 | 8.2 | 8.4 | 8.4 | 54.4 | 52 | 50.4 | 52.1 | 53.9 | | 42.1 | 2 | 1.9 | 2.2 | 2 | 2 | 2.1 | 2.3 | 2 | 2.4 |
| **NWAc7** | 7 | 6.8 | 7.5 | 7.6 | 7.1 | 6.6 | 7.4 | 6.9 | 7.3 | 7.3 | 43.7 | 42 | 42.6 | 42.5 | 44.5 | 41.4 | | 2 | 2.1 | 2.3 | 2.2 | 2 | 2.1 | 2.3 | 2.1 | 2.5 |
| **NEPc1** | 3 | 2.5 | 2.9 | 2.8 | 2.5 | 2 | 2.8 | 2.5 | 2.6 | 3.1 | 3.3 | 3.3 | 3.5 | 3.1 | 3.4 | 3 | 3.1 | | 36.1 | 36.4 | 35.5 | 35.7 | 36.4 | 37.9 | 40.6 | 37.9 |
| **NEPc2** | 2.9 | 2.7 | 3.1 | 2.9 | 2.5 | 2.9 | 2.7 | 2.5 | 3.1 | 3.2 | 3.4 | 3.4 | 3.4 | 2.8 | 3.4 | 2.8 | 3.2 | 34.5 | | 37.3 | 33.5 | 35.2 | 34.8 | 35.6 | 33.1 | 37.9 |
| **NEPc3** | 2.9 | 2.5 | 3 | 3 | 2.6 | 2.4 | 2.9 | 2.6 | 2.8 | 2.9 | 3.4 | 3.4 | 3.4 | 2.9 | 3.5 | 3.2 | 3.4 | 34.5 | 37 | | 34.1 | 34.4 | 40.6 | 37.5 | 32.3 | 58.3 |
| **NEPc4** | 3.1 | 3 | 3.4 | 3 | 3.1 | 2.3 | 3 | 2.7 | 3.2 | 3.1 | 3.5 | 3.5 | 3.8 | 3.1 | 3.5 | 3.2 | 3.6 | 37.5 | 37.1 | 38 | | 35.9 | 36.5 | 37.9 | 35.3 | 38.9 |
| **NEPc5** | 3.2 | 2.9 | 3.1 | 3.1 | 2.8 | 2.5 | 3.3 | 3 | 3.2 | 2.9 | 3.5 | 3.6 | 3.7 | 3 | 3.6 | 3.2 | 3.2 | 37.8 | 38.9 | 38.3 | 35.9 | | 36.6 | 37.8 | 33.8 | 39.3 |
| **NEPc6** | 2.7 | 2.4 | 3.1 | 2.9 | 2.6 | 2.3 | 2.6 | 2.7 | 2.5 | 2.9 | 3.3 | 3.3 | 3.3 | 2.8 | 3.2 | 3 | 3.1 | 34.6 | 34.6 | 40.6 | 32.8 | 32.9 | | 35.7 | 31.9 | 42.2 |
| **NEPc7** | 2.9 | 2.4 | 2.9 | 2.6 | 2.5 | 2.6 | 2.7 | 2.7 | 3 | 3.1 | 3.4 | 3.5 | 3.7 | 2.9 | 3.6 | 3.4 | 3.4 | 36 | 35.4 | 37.6 | 34.1 | 33.9 | 35.7 | | 32.4 | 38.8 |
| **NEPc8** | 3 | 2.5 | 3 | 2.8 | 2.5 | 2.3 | 3 | 2.5 | 3 | 3.2 | 3.5 | 3.8 | 3.6 | 3.5 | 3.7 | 3.2 | 3.3 | 41.3 | 35.2 | 34.7 | 34 | 32.5 | 34.2 | 34.7 | | 35.3 |
| **NEPc9** | 3.1 | 3 | 3.2 | 3.3 | 2.6 | 2.4 | 3.1 | 2.9 | 3.1 | 3 | 3.4 | 3.6 | 3.5 | 3.2 | 3.6 | 3.2 | 3.4 | 33.7 | 35.2 | 54.5 | 32.7 | 33 | 39.5 | 36.3 | 30.8 | |

**Supplementary table 6.** D-statistics testing for ancient admixture or “wrong tree” topology for the NEPp, NEPc and *Tursiops Aduncus;* NWAc, NWAp and *T. Aduncus*. The killer whale was used as the outgroup. H1 and H2 are the ingroup. nABBA os the total counts of ABBA patterns and nBABA the total counts of BABA patterns. D-stat is the results of nABBA-nBABA/nABBA+nBABA. A negative Dstat value indicate that H1 is closer to H3 than H2 while a positive value means that H2 is closer to H3 than H1 is. JackEst is an estimate of the abbababa statistic with bias correction. SE is the estimated m-delete blocked Jackknife Standard error of the estimate used to obtain the Z value. The Z score is used to determine the significance of the test. An absolute value of the Z score above 3 can be used as a significance threshold.

| H1 | H2 | H3 | nABBA | nBABA | Dstat | jackEst | SE | Z |
|---|---|---|---|---|---|---|---|---|
| NEPp | *Aduncus* | NEPc | 158613 | 1243205 | -0.77 | -0.77 | 0.00 | -296.64 |
| NEPc | Aduncus | NEPp | 159341 | 1243205 | -0.77 | -0.77 | 0.00 | -288.86 |
| NEPc | NEPp | *Aduncus* | 159341 | 158613 | 0.00 | 0.00 | 0.00 | 0.70 |
| NWAp | *Aduncus* | NWAc | 157778 | 1093409 | -0.75 | -0.75 | 0.00 | -302.00 |
| NWAc | *Aduncus* | NWAp | 194984 | 1093409 | -0.70 | -0.70 | 0.00 | -233.00 |
| NWAc | NWAp | *Aduncus* | 194984 | 157778 | 0.11 | 0.11 | 0.00 | 33.00 |

**Supplementary table 7.** List of the 45 genes associated with the SNPs under parallel selection (i.e. under both homogenising selection in the coastal populations and divergent between ecotypes showing both 1% FDR threshold, above which we consider the locus under selection), and putative functions

| Genes | Other gene names | Putative functions |
|---|---|---|
| **RPL10A** | | Encodes a ribosomal protein that is a component of the 60S subunit of the ribosomes |
| **TEAD3** | | Encodes the transcription factor TEF, which plays a role in the Hippo signaling pathway, a pathway involved in organ size control and tumor suppression |
| **GCM1** | | Encodes a transcription factor involved in the control of expression of placental growth factor (PGF) and other placenta-specific genes |
| **LOC101328860** | HLA-DQA2 | Encodes the HLA class II histocompatibility antigen, DQ alpha 2 chain, of immunologic importance |
| **LOC109552651** | ST3GAL1 | Encodes CMP-N-acetylneuraminate-beta-galactosamide-alpha-2,3-sialyltransferase 2-like. Cell type-specific expression of unique carbohydrate structures on cell surface glycoproteins and glycolipids provides information relevant to cell-cell interactions in developing and adult organisms. |
| **LPIN2** | | Encodes phosphatidate phosphatase LPIN2. Has a role in controlling the metabolism of fatty acids, and body composition, such as fat-mass ratio (61) |
| **LIAS** | | Encodes lipoyl synthase, involved in lipoic acid synthase, plays a central role in the antioxidant network |

| | | |
|---|---|---|
| **CAMK2D** | | Encodes a calcium/calmodulin-dependent protein kinase involved in the regulation of Ca(2+) homeostatis and excitation-contraction coupling (ECC). Targets also transcription factors and signaling molecules to regulate heart function |
| **LOC101323008** | MYH13-like | Encodes myosin-13 which is involved in muscle contraction |
| **LOC101324002** | MH3 | Encodes myosin-3, involved in heart contraction |
| **ADARB2** | ADAR3 | Encodes double-stranded RNA-specific editase B2. Has a role in RNA editing, and involve in cognitive, learning and memory abilities (62). |
| **MMP20** | | Encodes enamelysin, which has a role in dental enamel formation |
| **LOC109552895** | putative dimethylaniline monooxygenase [N-oxide-forming] 6 | COQ6 is a flavin-dependent monooxygenase needed for biosynthesis of coenzyme Q10. Coenzyme Q10 has a role as redox carrier in the mitochondrial respiratory chain and as a lipid-soluble antioxidant implicated in protection from cell damage by reactive oxygen species. |
| **LOC101318787** | G protein-coupled receptor 89A | Voltage dependent anion channel needed for acidification and functions of the Golgi apparatus |
| **ZNF697** | | Encodes the Zinc finger protein 697 and may be involved in transcriptional regulation |
| **DPYD** | | Encodes dihydropyrimidine dehydrogenase, involved in pyrimidine base degradation. Catalyzes the reduction of uracil and thymine. Also involved the degradation of the chemotherapeutic drug 5-fluorouracil. Involved in β-Alanine production, a putative neurotransmitter and a component of a number of coenzyme A and endogenous antioxidants found in the brain. |
| **STAR** | | Encode the steroidogenic Acute Regulatory Protein, involved in acute regulation of steroid hormone synthesis by enhancing the conversion of cholesterol into pregnenolone |
| **PALB2** | | Encodes Partner And Localizer Of BRCA2, involved in maintenance of genome stability, specifically the homologous recombination pathway for double-strand DNA repair |
| **MCM3AP** | | The protein encoded by this gene is a MCM3 binding protein, which are essential for the initiation of DNA replication |
| **RIOX2** | | Encodes the Ribosomal Oxygenase 2, which regulates immune responses and may play an important role in cell growth and survival |
| **GABRR3** | | The neurotransmitter gamma-aminobutyric acid (GABA) regulates synaptic transmission of neurons in the central nervous system. GABRR3 encodes one of three related subunits of the gene. Also has a potential role in aging and longevity. |

| | | |
|---|---|---|
| **CEP295** | | Encodes a centriole-enriched microtubule-binding protein with a role in elongating procentrioles after formation of the initiating cartwheel hub andposttranslational modification of centriolar microtubules |
| **TAF1D** | | TAF1D is part of the SL1 complex which has a role in RNA polymerase I transcription |
| **CUNH11orf54** | | Encodes ester hydrolase C11orf54 homolog which shows ester hydrolase activity on the substrate p-nitrophenyl acetate |
| **LOC101334760** | AGK – acylglycerol kinase | Encodes a lipid kinase involved in lipid and glycerolipid metabolism, catalyzes the formation of phosphatidic and lysophosphatidic acids |
| **CEP152** | | Encodes a centrosomal protein involved with centrosome function, has a role in cell shape, polarity, motility, and division |
| **LOC101331561** | GPR133, ADGRD1 adhesion G protein-coupled receptor D1 | Encodes a membrane-bound protein with long N termini containing multiple domains, possibly associated with adult height |
| **LOC101337631** | CYB5R4 | Encodes the cytochrome b5 reductase 4 involved in endoplasmic reticulum stress response pathway, protection of pancreatic beta-cells against oxidant stress and in in vitro reduction of cytochrome c, feericyanide and methemoglobin |
| **LOC101316550** | CCDC162P (coiled-coil domain-containing protein 162) | Transcripts from this locus encode truncated proteins, and may be involved in nonsense-mediated decay |
| **SERINC5** | | Encodes serine incorporator, which enhances the incorporation of serine into phosphatidylserine and sphingolipids, restrict infectivity of lentiviruses |
| **CCDC93** | | Protein coding gene, component of the CCC complex, which has a role in the regulation of endosomal recycling of surface proteins, including integrins, signaling receptor and channels |
| **ACER2** | | The ceramidase ACER2 hydrolyzes very long chain ceramides to generate sphingosine. Ceramides and sphingosine are bioactive lipids mediating cellular signaling pathways |
| **AAMP** | | Encodes a protein associated with angiogenesis and cell migration |
| **GPBAR1** | TGR5 | Encodes a member of the G protein-coupled receptor (GPCR) superfamily, which functions as a cell surface receptor for bile acids |

| | | |
|---|---|---|
| **FAM196A** | INSYN2A | Component of the protein machinery at the inhibitory synapses. This synaptic inhibition is central to the functioning of the central nervous system, it shapes and orchestrates the flow of information through neuronal networks to produce a precise neural code. |
| **PAOX** | | Encodes a Flavoenzyme which catalyzes the oxidation of N(1)-acetylspermine to spermidine and is involved in the polyamine back-conversion |
| **MTG1** | | Has a role in the regulation of the mitochondrial ribosome assembly and of translational activity, and shows mitochondrial GTPase activity |
| **RELN** | | Encodes the reelin protein, which has a role in the modulation of synaptic transmission in response to experience, learning and memory (63,64)[4]. |
| **RYR1** | | Encodes a ryanodine receptor found in skeletal muscle. The encoded protein functions as a calcium release channel in the sarcoplasmic reticulum, thereby have a key role in triggering muscle contraction and body movement. Can also mediate the release of calcium from intracellular stores in neurons. |
| **PATE2** | | PATE2 (Prostate And Testis Expressed 2) is a protein coding gene |
| **FEZ1** | | Encodes the Fasciculation and elongation protein zeta-1, which may be involved in axonal outgrowth |
| **FEZ2** | | Encodes the Fasciculation and elongation protein zeta-1, which is involved in axonal outgrowth and fasciculation |
| **CCDC57** | | Coiled-Coil Domain Containing 57 is a protein coding gene. Centrosomes function in key cellular processes ranging from cell division to cellular signaling |
| **SLC17A3** | | The protein encoded by this gene is a voltage-driven transporter that excretes intracellular urate and organic anions from the blood into renal tubule cells |
| **LOC101332501** | ZNF501 | Encodes the Zinc Finger Protein 501, which may be involved in transcriptional regulation |
